# An engineered bacterial symbiont maps micron-scale sugar gradients in the honeybee gut

**DOI:** 10.1101/2025.11.28.691108

**Authors:** Audam Chhun, Andrew Quinn, Alicia I. Pérez-Lorente, Théodora Steiner, Florian Zoppi, Thi Huong Giang Nguyen, Philipp Engel, Yolanda Schaerli

## Abstract

The honeybee gut microbiota plays a key role in shaping host health and susceptibility to disease. Yet, the nutrient environment it experiences within the gut remains poorly characterized. In particular, little is known about the spatial distribution of nutrients across the microbial community, as resolving such fine gradients in vivo has been technically challenging. Here, we engineer the native honeybee symbiont Snodgrassella alvi as a living biosensor to quantify the bioavailability of the dietary sugar arabinose within the gut. By expanding the genetic toolkit for S. alvi through chromosomal integration of high-burden genes and a suite of low-strength promoters, we achieve stable multi-gene expression without compromising host colonization. The resulting biosensor generates a specific, dose-dependent fluorescent response to arabinose in the living host, enabling visualization of sugar gradients across gut-associated bacterial biofilms at micron-scale resolution. Upon co-colonization with distinct Gilliamella species that differ in arabinose metabolism, the biosensor reported differential in vivo arabinose consumption, directly validating species-specific metabolic specialization within the host. Feeding bees with pollen further uncovered pronounced radial heterogeneity in the distribution of pollen- derived arabinose. These findings demonstrate how diet composition and microbial specialization generate fine-scale microenvironments within the gut. More broadly, this work establishes S. alvi as a genetically tractable platform for in situ biosensing, opening new avenues for dissecting metabolic interactions and nutrient distribution within living hosts.

## Introduction

The honeybee serves as a tractable model to study host-microbe interactions, as their gut harbors a simple, yet specialized bacterial community^1–3^. This microbiota exerts remarkable effects on its host, influencing nutrition^4,5^, pathogen resistance^6^, immunity^7^, detoxification^8^, behavior^9,10^ and cognitive development^11^. Chemical cues from both host and bacteria structure the community and govern these host-microbe and microbe-microbe interactions^12–14^. More specifically, it is the micron-scale distribution of those metabolites, rather than average levels, that dictates niche occupancy, cross-feeding and host phenotypes^15–17^. However, the gut chemical landscape remains challenging to spatially resolve in situ: the gastrointestinal tract is a highly dynamic, poorly accessible system with strong spatial heterogeneity.

Microsensors have profiled steep radial oxygen gradients and longitudinal pH variations at millimeter resolution in the bee gut^14^. In contrast, dietary carbohydrates, abundant from pollen and nectar, remain largely unmapped at comparable spatial scales, despite their central role in host health and disease^18,19^, and in the metabolic interplay of gut symbionts^4,20^. Current insight into localized sugar bioavailability is limited to demonstrations of carbohydrate breakdown in the stomach^21^, absorption in the midgut^22^ and to bulk chromatography- and mass spectrometry-based measurements that average signals across gut compartments^12–14^. Such approaches lose spatial context and ignore the important micro-scale gradients that steer microbial colonization. Tools capable of resolving carbohydrates in vivo with high spatial precision are therefore critical to advance our understanding of gut microbiota function within its native context.

Some progress toward spatial gut chemistry has been made in mice^23^ and some select invertebrates^24,25^ but have not been applied to insect models to the best of our knowledge. For instance, spatial host-microbe sequencing and correlative mass spectrometry imaging with fluorescence in situ hybridization have recently allowed unprecedented characterization of the mouse gut microenvironments^23,26,27^. Yet, these approaches often have limited sensitivity and resolution for small sugars, while requiring costly instrumentation and low throughput workflows. Another strategy is to deploy engineered symbionts as live biosensors to provide fine-scale, sensitive mapping of relevant metabolites directly in the gut^28,29^. Indeed, engineered bacterial biosensors have emerged as a versatile platform for interrogating host-associated environments, enabling the detection of metabolites, inflammatory markers and other transient environmental cues^30–36^, as well as recording transient signals through genetic memory circuits^37,38^. These studies have demonstrated the feasibility of programmable microbial sensors in vivo, but their application to insect microbiomes has remained largely unexplored. In honeybees, recent advances in the genetic engineering of gut commensals have established the core symbiont Snodgrassella alvi as a chassis for microbiome engineering and functional studies^39–41^. We previously engineered S. alvi to respond to the synthetic inducer IPTG, demonstrating the feasibility of cell-based sensing in this system and laying the groundwork for developing biosensors for ecologically relevant carbohydrates in the bee gut environment^42^.

Previous studies identified Gilliamella and Bifidobacterium as the principal degraders of complex carbohydrates in the bee gut^5^. Gilliamella apicola co-localizes with S. alvi in layered biofilms, suggesting metabolic crosstalk between these species^43^. Comparative genomic and metabolomic analysis indicate that they possess complementary metabolic abilities, whereby G. apicola deconstructs pollen glycans and ferments released sugars to organic acids that S. alvi consumes, in turn potentially allowing it to deplete oxygen and establish an anaerobic environment for the broader community^12,13,44^. Within this carbohydrate pool, arabinose is particularly interesting as high levels can be toxic to bees and only a subset of Gilliamella strains can catabolize it^5,45^. This sugar is commonly present in the bee diet, occurring in both nectar and pollen^46,47^. As such, arabinose represents a compelling target molecule for probing nutrient availability and interspecies metabolic coupling.

Here, we expand genetic engineering capacity in S. alvi by introducing synthetic promoters for fine control of complex circuits and a streamlined method for chromosomal integration of high-burden genes, enabling S. alvi to function as a living biosensor for the dietary sugar arabinose. The engineered strain shows a robust dose-dependent response in vitro and in vivo, and resolves micron-scale variation in arabinose within the gut. In co-colonization with Gilliamella strains that differ in carbohydrate metabolism, it reports arabinose consumption in a controlled diet. Leveraging its sensitivity, we uncover strong radial heterogeneity in the spatial distribution of arabinose derived from pollen degradation, resulting in nutrient gradients poised to structure the bee gut community. Together, these results establish engineered symbionts as powerful in situ reporters of metabolite bioavailability in the honeybee gut.

## Results

### Engineering the honeybee symbiont S. alvi as a biosensor for arabinose

Genetic analyses previously demonstrated that S. alvi lacks genes for import or utilization of carbohydrates as a carbon source^44^. To enable S. alvi to act as a biosensor for arabinose, we set out to genetically reprogram it to both import and respond to this sugar via the expression of a fluorescent protein (Fig. 1).

**Figure 1.**
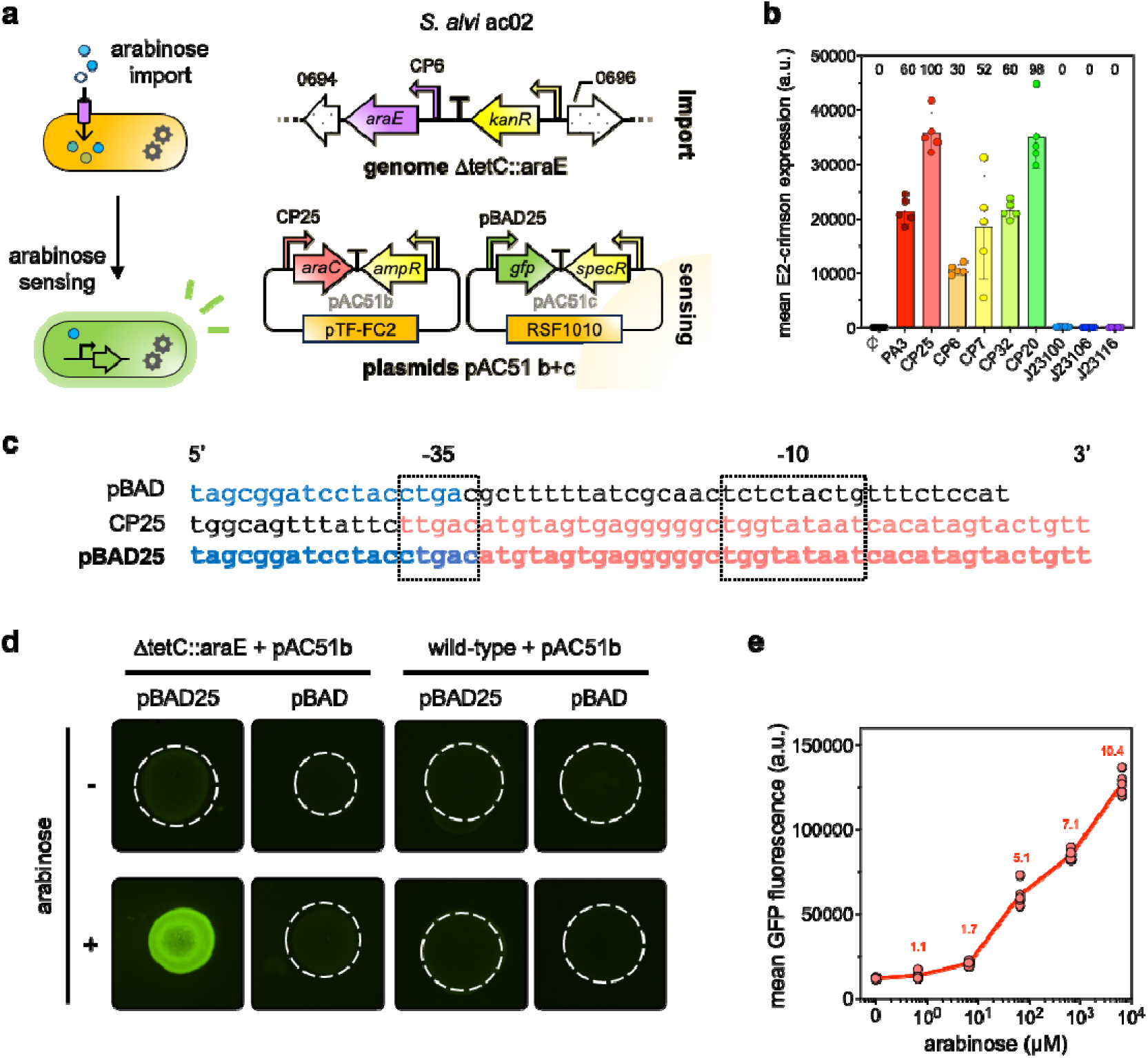
An engineered *S. alvi* strain detects and reports arabinose *in vitro* in a dose-dependent manner. **(a)** Overview of the genetic engineering carried out in the bee gut symbiont *S. alvi* to acquire arabinose import and sensing capabilities. The *araE* arabinose importer gene was integrated into the genome at the *tetC* locus, enabling arabinose import. Arrows filled with a dotted pattern represent neighboring genes flanking the insertion site and their gene names (SALWKB2_0694 and SALWKB2_0696) are shown above. The dual plasmid system pAC51, based on AraC-mediated regulation of the new synthetic arabinose-inducible promoter pBAD25, enabling arabinose sensing is also depicted. For clarity, certain regulatory elements have been omitted from the diagram. Complete annotations and sequence details are available in the Addgene repository. (b) Characterization of synthetic promoter in *S. alvi*. *S. alvi* cells were transformed with the plasmids pBTK570, pAC95, pAC82, pAC83, pAC84, pAC85, pAC86, pAC87, and pAC88; all bearing the E2-crimson gene driven by the promoters PA3, CP25, CP6, CP7, CP32, CP20, J23100, J23106 and J23116, respectively. As a negative control, *S. alvi* was also transformed with the plasmid pAC07, that does not include a fluorescent protein. PA3 and CP25 are commonly used promoters for engineering of S. alvi, displayed as reference here. The graph shows mean E2-crimson fluorescence values ± standard deviations of 5 biological replicates. Relative promoter activity (%, normalized to CP25) is indicated at the top. (c) The new arabinose-inducible pBAD25 promoter is a hybrid sequence comprised of the AraC-binding sites (i.e. O1L, O1R, I1 and I2) taken from the *E. coli-*derived pBAD promoter (in blue) and the core promoter region of CP25 (in red). Only partial sequences of pBAD and pBAD25 are shown for clarity, with 234 base pairs at the 5’ end omitted. RNA polymerase binding sites −35 and −10 are indicated by dotted rectangles. (d) Only S. alvi engineered with both heterologous expression of the chromosomal araE gene and the dual plasmid system based on the synthetic pBAD25 promoter respond to arabinose by expressing GFP. Top-view images of *S. alvi* bacterial cells grown on solid media for 3 days, supplemented with (+) or without (−) 13 mM arabinose. Dotted-line outlines the shape of non-fluorescent colonies. Photos were captured with a Fusion FX device (F-740 filter) using identical exposure times. (e) Strain *S. alvi* ac02 displays a dose-response to arabinose. The graph shows mean GFP fluorescence values from 5 biological replicates. Each replicate value was obtained from the average fluorescence of at least 9,000 bacteria measured by flow cytometry. Mean fluorescence fold-change relative to the uninduced condition is indicated in the same color. The data underlying this Figure can be found in the S1 Data file, sheets “Fig. 1b” and “Fig. 1e”.

To confer S. alvi arabinose import capability, we engineered cells to express AraE, a membrane protein responsible for arabinose transport in Escherichia coli^48^. Membrane proteins are difficult to express heterologously and prone to acquire mutations due to potential toxicity, requiring tight regulation of gene expression to maintain production at low levels^49,50^. In S. alvi, the few promoters that were available only result in high gene expression, while functional plasmid replication origins replicate in medium-to-high copy number, which together often cause instability in genetic circuits^40,42^. To address this limitation, we began by characterizing several synthetic promoters that, although previously published^51–53^, had not been evaluated in S. alvi. We identified the CP6 promoter as promising, with respectively 70% and 50% less activity than the CP25 and PA3 promoters typically used for engineering of this species^39,42^ (Fig. 1b). We then aimed to integrate the araE gene, driven by the CP6 promoter, into the *S. alvi* genome to minimize its copy number per cell. The previously reported example of targeted genome modification in *S. alvi* used a homologous recombination-based method to insert simple low-burden fluorescent reporters^41^. To achieve stable genomic integration of the membrane-protein encoding *araE* gene, we built upon this approach by developing a suite of standardized primers and plasmids containing different antibiotic resistance cassettes, allowing straightforward and versatile gene integration in *S. alvi* **(S1 Fig).** The constructs, referred to as suicide plasmids, are based on the conditional R6K origin of replication, which functions exclusively in strains expressing the *pir* gene. As a result, antibiotic-resistant S. alvi colonies can only arise through integration of the insert fragment, rather than replication of residual plasmids originating from the initial PCR reaction. We used these plasmids to pair the homology arms with an antibiotic cassette (kanR) and combined them in an overlap PCR with araE to generate a linear fragment for the *araE* knock-in. This strategy circumvented DNA replication in *E. coli*, thereby avoiding potential *araE*-associated toxicity and minimizing mutation risk (see Materials and Methods and **S1 Fig**). Ultimately, using this optimized genome engineering pipeline, we successfully integrated a genetically stable copy of the *araE* gene into the genome of *S. alvi* (**Fig. 1a**).

Next, to enable *S. alvi* to detect and respond to imported arabinose, we engineered a dual plasmid system in which AraC-mediated regulation drives GFP expression in response to the sugar **(Fig. 1a)**. In this design, *araC* and *gfp* are encoded on separate plasmids, as we previously found that multi-gene constructs are genetically unstable and prone to mutations in *S. alvi*^42^. As the *E. coli*-derived pBAD arabinose-inducible promoter^54^ is nonfunctional in this species, we designed pBAD25, a synthetic promoter variant functional in *S. alvi*, to control GFP expression **(Fig. 1a**; **Fig. 1c)**. *S. alvi* was able to respond to the presence of arabinose when engineered with both heterologous expression of the chromosomal *araE* gene and the synthetic pBAD25 promoter **(Fig. 1d)**. We then asked how key parameters, such as AraC and AraE expression levels, as well as plasmid copy number, could modulate the cells’ response to arabinose. For this, we constructed four *S. alvi* strains carrying variations of our genetic circuits and quantified their responses to a range of arabinose concentrations (**S2 Fig**). Each strain displayed a distinct sugar-response profile characterized by differences in dynamic range, sensitivity and basal leakiness (**S2b Fig**). Among these, the strain *S. alvi* ac02 exhibited the highest dynamic range and sensitivity, with fluorescence increasing up to 10.4-fold upon exposure to 6,661 μM arabinose relative to uninduced conditions, while maintaining low levels of basal leakiness (**Fig. 1e**). The detection limit of this strain was 6.7 μM arabinose, corresponding to a 1.7-fold increase in fluorescence signal (Welch’s t-test, t(4.6) = 14.8, p-value < 0.0001). Briefly, the strain carries a chromosomally integrated copy of the araE importer driven by the low-strength promoter CP6, while the dual plasmid system consists of a pTF-FC2-based plasmid bearing *araC* under constitutive expression from CP25 and a second RSF1010-based plasmid expressing GFP under the control of the inducible promoter pBAD25 (**Fig. 1a**). Based on these results, we concluded that *S. alvi* ac02 was a promising candidate to sense and respond in a dose-dependent manner to arabinose and pursued our work with this strain.

### In vitro characterization of the biosensor strain S. alvi ac02

Having engineered *S. alvi* to detect and respond to arabinose, we next sought to characterize *in vitro* the strain’s properties relevant to its function as a gut biosensor. We examined (*i*) the specificity of the response to arabinose; (*ii*) the cell sensitivity to ecologically relevant sugar concentrations, and (*iii*) the stability of the genetic modifications in the absence of selection pressure.

The engineered strain needed to distinguish arabinose from structurally related sugars. The honeybee diet is indeed remarkably rich in carbohydrates, including the pentose monosaccharides xylose, ribose and arabinose^45^. These sugars are stereoisomers that differ only in the spatial arrangement of a hydroxyl group. To test if our biosensor responds specifically to arabinose, we grew cells in media supplemented with 0.7 mM of either ribose, xylose or arabinose, as well as equimolar mixtures of arabinose with either ribose or xylose, and measured their fluorescence levels (**Fig. 2a**). The strain responded selectively to arabinose, with a 4.5-fold increase in GFP signal, while exposure to ribose or xylose resulted in fluorescence levels comparable to the uninduced condition. Consistent with the well-documented specificity of the AraC-pBAD regulatory system for L-arabinose over related pentoses^54–56^, these results indicate that the engineered strain S. alvi ac02 provides a specific and reliable readout of arabinose in the presence of structurally related carbohydrates that are found in the bee diet.

**Figure 2.**
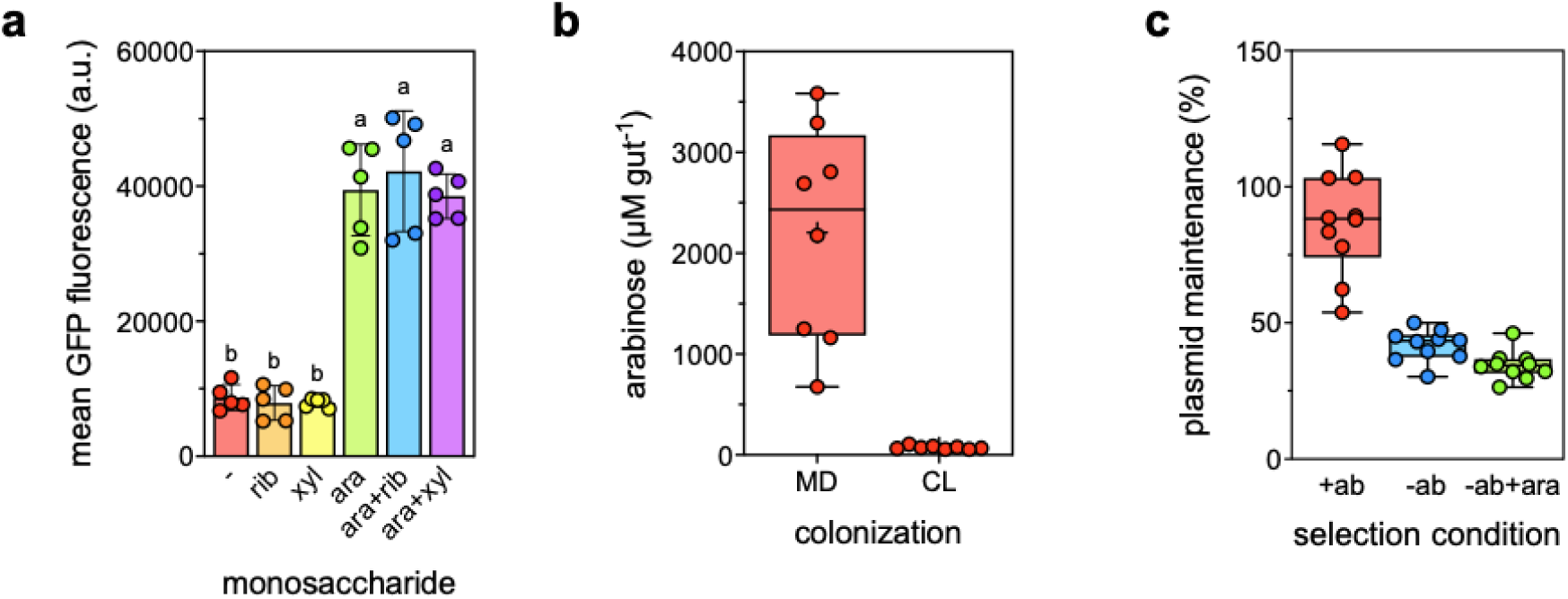
*In vitro* characterization of the biosensor *S. alvi* ac02 properties. **(a)** *S. alvi* ac02 responds specifically to arabinose. Cells were grown for 3 days in TSB alone (−) or in TSB supplemented with 0.7 mM of ribose (rib), xylose (xyl), arabinose (ara) or equimolar mixtures of ribose/arabinose (ara+rib) and xylose/arabinose (ara+xyl), each at 0.7 mM. Bar plots show mean of GFP fluorescence ± standard deviation of 5 biological replicates. Each replicate value was obtained from the average fluorescence of at least 9,000 bacteria measured by flow cytometry. Different letters indicate significant differences between sugars with adjusted *p*-value < 0.0001 (Tukey HSD test). **(b)** Pollen-derived arabinose concentrations in bee guts. Bees were fed with sugar water and pollen. They were either microbiota-deprived (MD) or colonized (CL) for 7 days prior to dissection and arabinose quantification. Box plots show median arabinose concentrations measured by GC-MS and normalized with the average volume of a bee gut (n = 8). **(c)** Maintenance of arabinose-inducible plasmids pAC51b and pAC51c in *S. alvi* ac02 in the absence of antibiotic selection. Each replicate represents a 3-day liquid culture grown in TSB with ampicillin and spectinomycin (+ab), TSB alone (-ab) or TSB with arabinose (-ab+ara). Plasmid maintenance was calculated as the fraction of CFUs growing on TSA with both antibiotics (*i.e.* cells bearing the two plasmids pAC51b and pAC51c) relative to total CFUs on non-selective TSA. Box plots show median plasmid maintenance of 10 biological replicates. The data underlying this Figure can be found in the S1 Data file, sheets “Fig. 2a”, “Fig. 2b” and “Fig. 2c”.

We then asked whether the biosensor could respond to sugar concentrations representative of those present in the honeybee gut after pollen digestion. To determine these levels, we dissected guts from pollen-fed, microbiota-deprived bees (MD) or individuals colonized by a defined bacterial community of 25 isolates (CL, see details in the methods), homogenized them in 500 μL sterile PBS, and analyzed them by mass spectrometry (**S3a Fig**). To convert mass spectrometry values into realistic in vivo concentrations, we estimated the average honeybee gut volume, by measuring the mass loss of water after desiccation, to be 19.1 ± 4.7 μl (see Materials and Methods and **S3b Fig**). Using this normalization for the gut volume, arabinose concentrations averaged 2,204.4 ± 1,071.1 μM in MD bee guts, and 72.2 ± 16.4 μM in CL bee guts (n = 8; Fig. 2b). This range falls within the detection window of the engineered S. alvi strain ac02 (**Fig. 1e**), suggesting it can report variations in arabinose concentrations of both extremes: high concentrations in the absence of microbial metabolism (*i.e*. MD bees) and reduced, levels when a gut community consumes arabinose (*i.e*. CL bees).

Next, we examined the stability of the genetic circuits in the absence of antibiotics. While part of the biosensor design was chromosomally integrated, we hypothesized that plasmid-borne elements might be lost under non-selective conditions, as expected in the natural gut environment^42^. To test this, we grew S. alvi ac02 in liquid tryptic soy broth (TSB) medium with antibiotics (+ab), without antibiotics (-ab), or without antibiotics and arabinose supplementation (-ab+ara; i.e. with the circuit activated) and then isolated them onto selective and non-selective solid media. We quantified plasmid maintenance as the fraction of bacteria retaining both plasmids (**Fig. 2c**). Under antibiotic pressure, plasmids were stably maintained whereas in its absence maintenance declined to 41.7 ± 5.8 % (-ab) and 34.4 ± 5.3 % (-ab+ara). We found that loss was specific to the araC-bearing plasmid pAC51b while the gfp-bearing plasmid pAC51c remained stable (**S4a Fig**). Despite this reduction, the absolute number of plasmid-bearing cells remained consequential even when they were exposed to arabinose, with an average of 4.3 x 10^7^ ± 5.3 x 10^6^ CFU ml^-1^ (n = 10; **S4b**), suggesting that enough bacterial biosensors should remain functional in the gut environment. To ensure that plasmid dynamics were not associated with a growth defect, we also assessed whether the biosensor constructs affect bacterial proliferation. We quantified CFU counts of *S. alvi* cells carrying the dual-plasmid system (pAC51b/c; +) or lacking the plasmids (−) following growth to stationary phase in the presence of different arabinose concentrations (**S4c Fig**). We detected no significant differences in cell biomass either across arabinose concentrations or between plasmid-containing and plasmid-free strains (two-way ANOVA, effect of plasmids: F(1, 24) = 2.68, p-value = 0.114; effect of arabinose: F(5, 24) = 0.48, p- value = 0.785), indicating that the biosensor system does not impose a measurable growth burden under the tested conditions.

### Biosensor-based mapping of arabinose distribution in the honeybee gut

Building on the in *vitro* demonstration of the biosensor’s potential, we next evaluated its performance *in vivo*. Specifically, we focused on whether the engineered strain could establish in the honeybee gut, how long it could persist and whether it could sense arabinose *in situ* and provide spatial information on sugar availability.

First, we assessed the biosensor ability to colonize the honeybee gut. For this, microbiota-deprived bees were either mono-colonized with the engineered strain *S. alvi* ac02 and fed sugar water with (+ab) or without antibiotics (-ab), or co-colonized with the biosensor and a native undefined community derived from gut homogenate that includes wild-type *S. alvi* (-ab+gc; **Fig. 3a**). We monitored colonization dynamics by quantifying bacterial loads in fecal samples over time. Results showed that even in the absence of antibiotic selection, colonization was highly successful: 100% of mono-colonized bees (25 of 25 bees) and 95.6% of co-colonized bees (22 of 23 bees) harbored the biosensor after 7 days, with mean bacterial loads of 4.3 x 10^5^ ± 5.7 x 10^5^ CFU μl^-1^ and 6.0 x 10^3^ ± 9.2 x 10^3^ CFU μl^-1^, respectively. However, biosensor abundance declined by day 14, most notably in co-colonized bees, whereas mono-colonized individuals remained colonized albeit to lower bacterial loads (5.6 x 10^2^ ± 2.3 x 10^3^ CFU μl^-1^, 68.2% of bee colonized). Based on these observations, all subsequent measurements were performed 7 days post-colonization.

**Figure 3.**
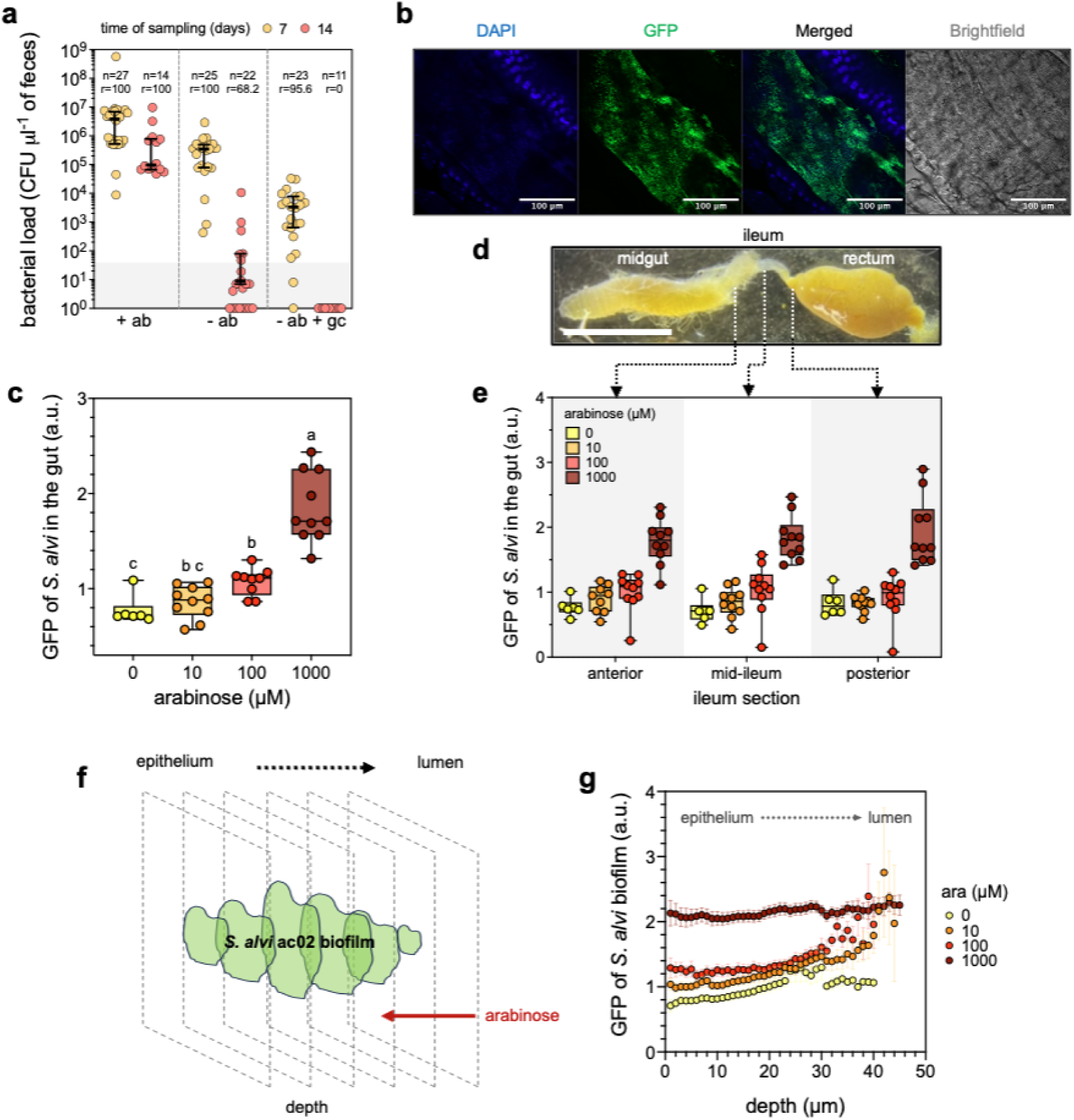
Engineered biosensor enables high-resolution mapping of arabinose in the honeybee gut. **(a)** Bacterial load of engineered *S. alvi* ac02 cells isolated from bee feces collected 7 (orange) and 14 (red) days after colonization. Bees were either mono-colonized with the biosensor strain and fed sugar water supplemented with (+ab) or without (−ab) antibiotics; or co-colonized with the biosensor strain and the native gut community (-ab+gc). Bees for which engineered bacteria were not detected in the feces were considered non-colonized. The rate of colonized bees (r) and the total number of individuals sampled (n) are indicated. For colonized bees, colored horizontal bars show median values with interquartile ranges. **(b)** A confocal microscopy image of a honeybee gut colonized by engineered *S. alvi* ac02 in the ileum region. The bee was fed sugar water without arabinose supplementation. A bacterial biofilm within a gut crypt is shown. The blue channel depicts DAPI staining of host and bacterial DNA and the green channel shows basal GFP fluorescence from the engineered strain. **(c)** *S. alvi* ac02 responds in a dose-dependent manner to arabinose *in vivo.* Box plot shows median value of GFP fluorescence of bacterial biofilms imaged from the gut of bees fed sugar water supplemented with indicated concentrations of arabinose (n = 10). Gut dissection was performed 7 days post colonization. Each fluorescence value corresponds to the average intensity measured from 3 gut sections. Different letters indicate significant differences between arabinose concentrations with adjusted *p*-value < 0.1 (Tukey HSD test) (d) Stereo microscopy image of a honeybee gut. The different gut regions are indicated and separated by white dotted lines. Scale bar represents 5 mm. (e) Sugar derived from a simple diet is homogeneously distributed along the longitudinal axis of the honeybee gut. Box plots show median value of GFP fluorescence of *S. alvi* ac02 biofilms found in different gut regions sourced from bees fed sugar water supplemented with varying concentrations of arabinose (n = 10). Gut dissection was performed 7 days post colonization. The regions considered were the anterior (ant) region adjoining the midgut, the mid-ileum (mid) and the posterior (post) region adjoining the rectum. Data was analyzed using a two-way ANOVA. There was no significant effect of ileum section (F(2, 95) = 0.03, p-value = 0.972), a significant effect of arabinose concentration (F(3, 95) = 70.8, p-value < 0.0001), and no significant interaction between ileum section and arabinose concentration (F(6, 95) = 0.47, p-value = 0.832). (f) Schematic of the microscopy rationale for sugar distribution analysis across bacterial biofilm. (g) Sugar derived from a simple diet distributes along a gradient across the depth of *S. alvi* ac02 biofilm. Graph shows mean GFP fluorescence ± SEM of the biosensor according to the biofilm depth. For each bee (n = 10), GFP signal was first averaged over the three ileum regions. The graph displays the average of these per-bee means. Only fluorescence values for which at least two replicates could be measured for a given depth are reported. The data underlying this Figure can be found in the S1 Data file, sheets “Fig. 3a”, “Fig. 3c”, “Fig. 3e” and “Fig. 3g”.

To determine whether the biosensor remained genetically stable during gut colonization, we additionally quantified plasmid maintenance *in vivo* by plating fecal samples from bees fed with or without antibiotic supplementation on selective media. Consistent with our *in vitro* observations, the *gfp*-bearing plasmid pAC51c was maintained at high levels throughout the two-week experiment (75.3 ± 23.5 % without antibiotic selection), whereas maintenance of the *araC*- bearing plasmid pAC51b declined in the absence of antibiotic selection (**S5a Fig**). Notably, pAC51b retention was similar *in vitro* and *in vivo*, averaging 37.8 ± 6.9 % and 33.5 ± 27.6 %, respectively (Mann-Whitney U test, p-value = 0.198). To assess whether plasmid-bearing cells remained functional, we further tested arabinose-induced GFP expression in colonies recovered from the same colonized bees at day 7. Cells isolated from bees fed both antibiotic-supplemented and normal sugar water retained robust inducibility (**S5b Fig**), indicating that the biosensor remained functional throughout gut colonization over the time frame examined.

Having confirmed successful colonization, persistence of the genetic circuitry and maintenance of biosensor functionality in the honeybee gut, we next investigated the biosensor capacity to sense arabinose in vivo. To quantify the *in vivo* response of our biosensor, we mono-colonized bees with *S. alvi* ac02 and fed them sugar water containing varying concentrations of arabinose. After 7 days, we dissected guts and imaged them by confocal microscopy. A low basal level of GFP expression in the absence of arabinose enabled straightforward identification of the biosensor bacteria, even without induction. We confirmed that the engineered strain successfully colonized the honeybee gut and formed biofilms that could be readily visualized (**Fig. 3b**). We then quantified fluorescence intensity with a custom FIJI pipeline that automatically defined the biofilm region. The biosensor strain displayed a clear dose-dependent response to arabinose, with fluorescence increasing significantly with higher arabinose concentrations (**Fig. 3c**).

With colonization and dose-dependent sensing demonstrated, we next examined whether the biosensor could resolve spatial variations in sugar availability both longitudinally (*i.e*. along the length of the gut) and transversally (*i.e*. across the gut). To assess longitudinal variation, fluorescence was measured in three distinct regions of the ileum: the anterior region adjoining the midgut, the mid-ileum and the posterior region adjoining the rectum (**Fig. 3d**). The biosensor reported fluorescence intensities consistent with the arabinose concentrations fed to the host but exhibited no longitudinal variation, with similar fluorescence levels across all three regions for a given sugar condition (**Fig. 3e**; two-way ANOVA, effect of ileum section not significant: F(2, 95) = 0.03, *p*-value = 0.972). These results indicate that arabinose is homogeneously distributed along the ileum’s length.

Having established longitudinal uniformity, we next asked whether arabinose availability varied transversally (*i.e*. across the bacterial biofilm on the host epithelium). We hypothesized that micro-scale nutrient gradients exist in these biofilms, because cells located closer to the lumen typically experience higher nutrient access, while those deeper in the biofilm encounter reduced concentration^57,58^. To test whether our biosensor could resolve such fine-scale gradients, we performed confocal imaging with 1 μm depth increments, tracking biofilms of the engineered cells from their epithelial attachment sites to the gut lumen (**Fig. 3f, S6 Fig**). In bees fed no arabinose, fluorescence remained uniform across the biofilm depth, whereas a progressive increase in fluorescence toward the lumen was observed in bees fed 10 and 100 μM arabinose, revealing a sugar gradient within the biofilm (**Fig. 3g**). Notably, in bees fed 1,000 μM arabinose, the fluorescence signal was evenly distributed throughout the biofilm, consistent with more uniform substrate availability across the biofilm at increased concentrations. Together, these results validate the *S. alvi* biosensor as an in vivo reporter capable of resolving transversal sugar gradients within gut biofilms, while revealing longitudinal uniformity along the ileum.

### *In vivo* mapping of pollen-derived arabinose gradients in bees colonized with *Gilliamella*

After establishing that engineered *S. alvi* resolves *in vivo* arabinose gradients under a simplified diet, we used it to probe the spatial distribution of pollen-derived sugar during co-colonization with members of the genus *Gilliamella*.

Genomic analysis suggests divergent arabinose metabolism between the strains *G. apis* ESL169 and *G. apicola* ESL309. The former lacks genes involved in arabinose import and catabolism, whereas the latter has the gene to use this sugar as a carbon source. To visualize strains in situ, both *Gilliamella* strains were tagged with a plasmid encoding for the fluorescent reporter E2-crimson, allowing to discriminate the cells from GFP-expressing S. alvi. We co-colonized honeybees with the *S. alvi* biosensor and either *G. apis* ESL169 or *G. apicola* ESL309 (**Fig. 4a**), housed the bees in cup-cages and fed them one of three diets: *(i)* simple sucrose sugar water (SW), *(ii)* SW supplemented with arabinose or *(iii)* SW and pollen ad libitum. As pollen exhibits autofluorescence primarily in the red spectrum (**S7 Fig**), which may interfere with E2- crimson signal detection, pollen was removed from cages after 5 days. After 7 days, we dissected the guts and imaged them by confocal microscopy. Image analysis confirmed that both the *S. alvi* engineered strain and *Gilliamella* could successfully colonize the honeybee gut and co-localized in the ileum as layered biofilms (**Fig. 4b**).

**Figure 4.**
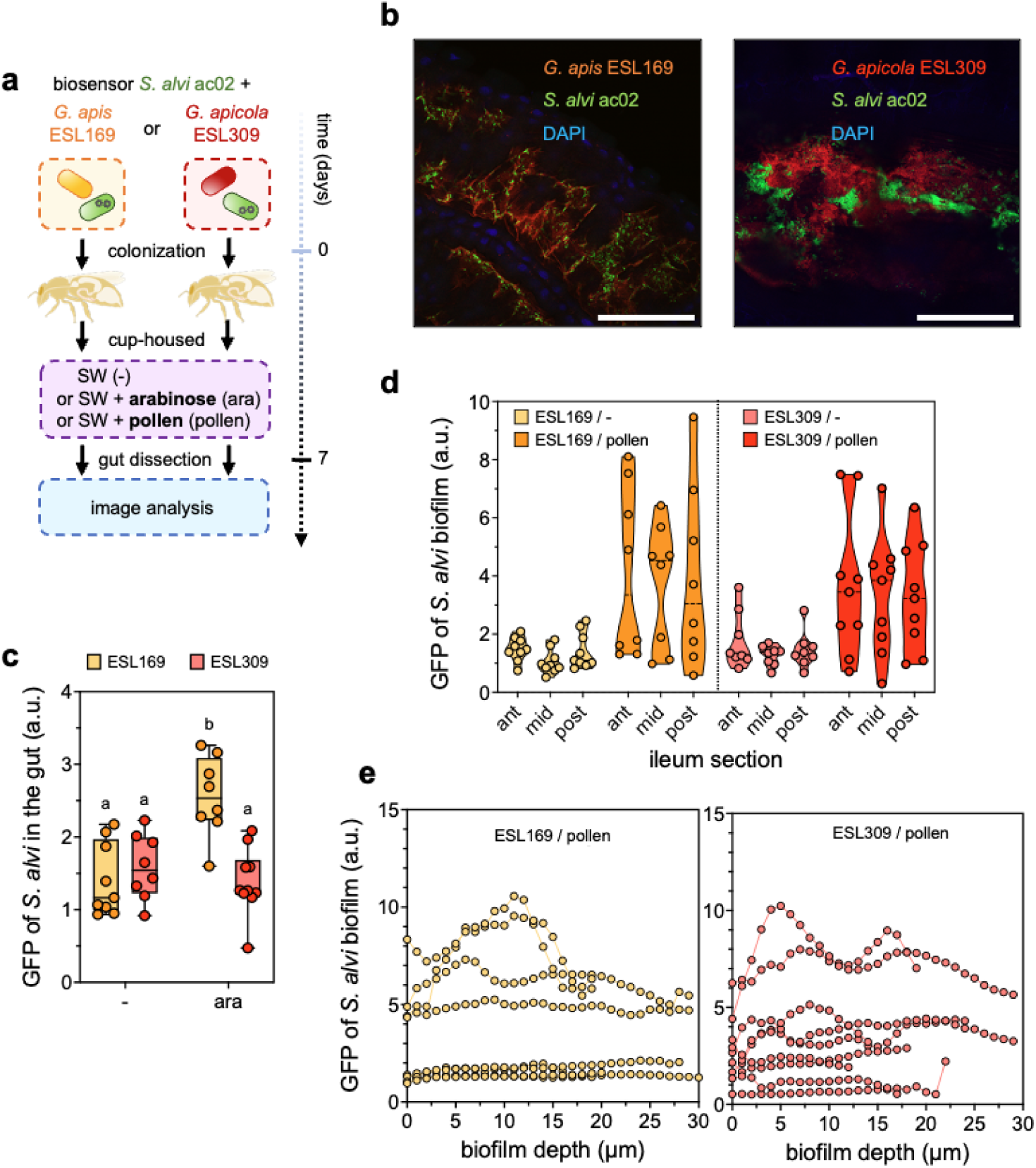
Biosensor mapping reveals heterogenous arabinose gradients from pollen degradation. **(a)** Schematic outline of the *in vivo* experiment workflow. All gut dissection and image analysis were performed 7 days post-colonization. Bee illustration reproduced from ref. 42. **(b)** Confocal microscopy images of a honeybee ileum co-colonized by *S. alvi* ac02 and *G. apis* ESL169 (left) or *G. apicola* ESL309 (right). The blue channel depicts DAPI staining of host and bacterial DNA. The green channel shows induction of GFP fluorescence in the engineered *S. alvi* strain, triggered by pollen-derived arabinose. Red channel shows E2-crimson fluorescence from the *Gilliamella* strains transformed with the pAC09 plasmid. Scale bar represents 100 μm. **(c)** *G. apicola* ESL309 catabolizes arabinose whereas *G. apis* ESL169 does not. Box plot shows median value of GFP fluorescence of *S. alvi* biofilms imaged from the gut of bees fed sugar water (−) or sugar water supplemented with 210 μM arabinose (ara). Each fluorescence value corresponds to the overall mean intensity averaged from 3 sections (anterior, mid-ileum, posterior) of one gut and across the depth of the bacterial biofilm for each section. Different letters indicate significant differences in fluorescence intensities across colonization and diet conditions, with adjusted *p*-value < 0.05 (Tukey HSD test). **(d)** Distribution of pollen-derived arabinose is longitudinally uniform along the ileum. Truncated violin plot shows median value of GFP fluorescence quantified from *S. alvi* biofilms imaged in the gut of bees fed sugar water (−) or sugar water and pollen *ad libitum* (pollen). Each fluorescence value corresponds to the mean intensity averaged across the depth of the bacterial biofilm for each gut section. Data was analysed using a three-way ANOVA. There was no significant effect of ileum section (F(2, 90) = 0.77, *p*- value = 0.468), no significant effect of species (F(1, 90) = 0.07, *p*-value = 0.796) and a significant effect of food (F(1, 90) = 12.77, *p*-value = 0.001). **(e)** Arabinose derived from pollen breakdown distributes with high heterogeneity along a radial gradient across the depth of *S. alvi* ac02 biofilm when co-colonized with *G. apis* ESL169 (left) or *G. apicola* ESL309 (right). Graphs show the mean GFP fluorescence of the biosensor according to the biofilm depth of individual bees. Data corresponding to the anterior ileum region are displayed only for clarity. Complete data are provided in S9 Fig. The data underlying this Figure can be found in the S1 Data file, sheets “Fig. 4c”, “Fig. 4d” and “Fig. 4e”.

We next aimed to confirm the metabolic abilities of the *Gilliamella* strains with regards to arabinose *in vivo*. For this, we quantified *S. alvi* fluorescence across colonization states and diets. The biosensor was induced only in arabinose-fed bees co-colonized with *G. apis* ESL169 (Fig. 4c), consistent with arabinose accumulation in the gut. In contrast, no induction was observed with *G. apicola* ESL309, indicating depletion of arabinose by the strain. These results show that the biosensor can report on *in vivo* arabinose consumption and thus validate the divergent sugar metabolisms of the two *Gilliamella* strains tested. Additionally, our observations provide evidence that species-level differences in *Gilliamella* can influence sugar availability in the gut.

Having demonstrated the efficacy of the biosensor to report on arabinose metabolism in co-colonized bacteria, we then leveraged it to map locally the spatial distribution of pollen-derived arabinose. Pollen-based diet resulted in significant induction of the biosensor, indicating the accumulation of detectable levels of arabinose in the ileum, which were notably independent of the *Gilliamella* species present (**Fig. 4d**; three-way ANOVA, effect of species not significant: F(1, 90) = 0.07, *p*-value = 0.796). Also, fluorescence measurements of *S. alvi* along the ileum’s length revealed highly heterogenous arabinose concentrations between bees, although homogeneously distributed longitudinally (**Fig. 4d**; three-way ANOVA, effect of ileum section not significant: F(2, 90) = 0.77, p-value = 0.468) and overall consistent within each individual (**S8 Fig**). By contrast, depth-resolved quantification across *S. alvi* biofilms showed pronounced radial heterogeneity, with arabinose availability varying sharply over microns within a same biofilm and across individuals (**Fig. 4e; S9 Fig**). Thus, while arabinose is longitudinally uniform along the ileum, pollen degradation by host enzymes and bacterial metabolism generates patchy, micron-scale nutrients niches within biofilms that differ between bees. These results establish engineered symbionts as sensitive, spatially precise reporters of sugar bioavailability in vivo, enabling mapping of complex gut chemical microenvironments.

## Discussion

Here, we engineered the native honeybee gut symbiont *S. alvi* as a living biosensor that reports the bioavailability of the dietary sugar arabinose within the gut. By expanding the genetic toolkit for *S. alvi*, we reprogrammed the bacterium to produce a specific, dose-dependent response to arabinose both *in vitro* and *in vivo*, resolving micron-scale nutrient variation across the gut environment. Using this system, we reveal pronounced spatial heterogeneity in arabinose distribution derived from pollen degradation, uncovering nutrient gradients that likely influence the spatial organization and dynamics of the bee gut microbiota. When co-colonized with distinct *Gilliamella* species, the biosensor reported species-specific arabinose depletion under defined dietary conditions, exposing differences in their carbohydrate metabolism.

Our work confirms *S. alvi* as a genetically accessible model system for functional engineering of the bee gut microbiota. Although this species has been engineered before, existing genetic tools could not accommodate complex circuit designs, as only a few high-expression promoters were available^40^. The limited diversity of promoters forced the use of repeated identical sequences across multi-gene constructs and their strong activity imposed substantial metabolic burden, both factors leading to recombinations and mutations^42^. By combining chromosomal integration with a suite of low-strength promoters, we overcome these limitations, achieving stable and predictable multi-gene expression in *S. alvi* without compromising its capacity to colonize the gut, albeit within a restricted but experimentally relevant time window.

While our data indicate that the biosensor remains functional within a 7-day experimental window, the genetic stability of the construct over longer time scales was not directly assessed, and the potential accumulation of non-functional mutations or plasmid loss at later time points may influence its performance for longer term measurements. Future work could address this limitation by quantifying functional stability over more extended time periods and by implementing more stable genetic designs. The number of well-characterized and reliably neutral genomic integration sites (*i.e*. landing pads) in *S. alvi* remains, however, limited. Expanding the repertoire of defined landing pads would facilitate the chromosomal encoding of increasingly complex genetic circuits, reducing reliance on plasmids and further enhancing the long-term stability of engineered strains.

Nevertheless, the expanded genetic control presented here enhances the tractability of *S. alvi* and broadens its potential as a chassis for *in vivo* studies and microbiome engineering. Building on our previous work with synthetic inducers, the expanded toolkit now enables the implementation of more complex functionalities for nutrient biosensing within the gut environment, while providing a foundation for developing additional biosensors tailored to the diverse chemical signals encountered by symbionts during colonization.

Developing biosensors in non-model symbionts presents challenges beyond transferring an existing sensing module into a new host. In *S. alvi*, implementation of the arabinose sensor required functional uptake of the target molecule, development of genetic parts with appropriate expression levels, reliable chromosomal integration, and construction of a synthetic promoter that responds to arabinose in this organism. More generally, sensor performance in a native symbiont depends on the compatibility of its components with the bacterium’s genetic and physiological constraints, including endogenous regulatory and metabolic networks, genetic stability, and its physiological state during gut colonization. The genetic framework developed here addresses some of these challenges and provide a basis for future development, although each new sensing system will require specific optimization and validation to achieve robust *in vivo* performance, particularly when target molecules differ in their uptake or regulatory mechanisms.

A further challenge is matching sensor performance to the concentration range encountered in the native gut environment. Here, our arabinose biosensor can detect a biologically relevant range of arabinose concentrations with sufficient sensitivity. For applications requiring detection of lower concentrations, however, future iterations of the biosensor could increase sensitivity, for example, by incorporating a genetic amplification circuit^59^. At the same time, such enhancements would need to be balanced with maintaining stable and reliable performance *in vivo*.

The ability of an engineered honeybee symbiont to sense and report sugar *in situ* provides a compelling demonstration of how synthetic approaches can provide new insights into previously inaccessible features of gut chemical landscapes. Past efforts to characterize nutrient distribution in animal microbiomes have largely relied on bulk chemical measurements (*i.e*. feces or homogenized gut tissues) or indirect inference using techniques like metagenomics^60^ or isotope tracing^61^. These methods fail to capture fine-scale spatial and temporal heterogeneity. In contrast, our biosensor strain enables metabolite detection within the native gut environment, preserving both host and microbial context. Recent advances in spatial metabolomics, especially mass spectrometry imaging, have also begun to probe metabolite distribution in situ, typically achieving resolutions in the 3-10 μm range with commercial setups^17,24^. Notably, we could here resolve sugar variation within bacterial biofilms at the 1 μm scale. By engineering biosensing capabilities into a native symbiont with this spatial precision, our approach achieves finely resolved nutrient detection that remains ecologically relevant within its natural niche.

This resolution proved particularly powerful under a simplified diet of arabinose-supplemented sugar water, where it revealed a transversal arabinose gradient across bacterial biofilms, with a higher concentration detected closer to the gut lumen. It also enabled the discrimination of species-specific arabinose metabolism among co-colonizing *Gilliamella* species. However, these metabolic differences were no longer detectable under a more complex, pollen-based diet. Instead, the biosensor exposed pronounced radial heterogeneity in arabinose distribution, likely arising from the continued degradation by host enzymes and bacteria of partially digested pollen grains that can be observed embedded within bacterial biofilms. These residual pollen fragments may act as localized nutrient-release sites, generating patchy, micron-scale variations in sugar availability. The strong heterogeneity in pollen-derived arabinose concentrations observed between bees might also reflect temporal and behavioral effects. For instance, differences in how recently individual bees consumed pollen prior to gut dissection or in how strongly they favored sugar water versus pollen feeding. Regardless of its exact origin, this heterogeneity was independent of the Gilliamella species present. This is likely due to the high overall arabinose content of pollen, where small-scale depletion by individual species was insufficient to alter the biosensor’s response. Alternatively, the presence of pollen-derived metabolites or the limited competition for substrates inherent to this two-member co-colonization may modulate *Gilliamella* metabolism towards reduced arabinose utilization. Nonetheless, these observations demonstrate how diet complexity and pollen breakdown can generate heterogeneous nutrient micro niches within the gut, which can shape microbial interactions at fine spatial scales.

Our work establishes a framework for probing the spatial distribution of nutrients in the honeybee gut at an unprecedented resolution, using arabinose as a model substrate. However, pollen degradation releases a much broader spectrum of molecules that includes other carbohydrates, proteins, lipids, amino acids and vitamins^47^. They likely form overlapping chemical gradients shaping community structures and interactions. Indeed, distinct members of the honeybee gut microbiota occupy different metabolic and spatial niches. *Lactobacillus* species, for instance, can utilize sugar alcohols and flavonoids, whereas *Bifidobacteria* dedicate a large fraction of their catabolism to the breakdown of hemicellulose and other complex polysaccharides^5,12,20^. Extending biosensing to these compounds, or simply to other sugars, would provide a more comprehensive view of how diet composition and colonization state structures the gut’s metabolic environment. While arabinose has been reported to exert toxicity to honeybees at concentrations more than two orders of magnitude higher than those measured in our study^62^, other sugars like mannose are known to adversely affect bees due to inefficient metabolism and accumulation of mannose-6-phosphate^63^. Mapping the distribution and bioavailability of such sugars *in situ* could therefore yield valuable insights into how specific dietary components impact gut function and host physiology. In addition, the sugar composition of bee pollen varies greatly depending on plant origin^64^, suggesting that the nutrient landscape experienced by bees feeding on distinct wildflower mixes or single-crop diets may differ substantially. Leveraging our biosensor to monitor how these dietary differences shape metabolite availability would help clarify how food sources could influence the gut environment. Overall, achieving this will require further expansion of the *S. alvi* genetic toolkit, with sensors responsive to a broader range of metabolites and genetic modules supporting more complex circuit designs. In parallel, implementing these biosensors in multi-species or fully reconstituted communities will be essential to capture the metabolic interplay that occurs under natural colonization conditions. Together, these advances would enable high-resolution, community-level mapping of nutrient fluxes and functional specialization within the honeybee gut ecosystem.

Finally, our findings highlight the potential of synthetic biology to bridge microbial ecology and functional genomics in natural host systems. By combining genetic engineering with *in situ* readouts, this work demonstrates how non-model symbionts can be transformed into quantitative tools to study metabolism for microbiome research. Expanding such approaches across hosts and microbial taxa will require continued development of genetic toolkits for undomesticated bacteria, an effort that remains a major bottleneck for applications beyond traditional laboratory organisms. As these capabilities mature, engineered symbionts will open new avenues to uncover the complex chemical dialogues that underpin host-microbe symbiosis.

## Materials and methods

### Bacterial strains and culture conditions

The honeybee gut symbionts *Snodgrassella alvi* wkB2^T^ (ATTC No. BAA-2449), the engineered biosensor strain *S. alvi* ac02, *Gilliamella apicola* ESL309 and *Gilliamella apis* ESL169 were routinely cultured in Tryptic Soy Broth (TSB, Bacto BD). Bacteria were grown at 34 under microaerophilic conditions within a 5 % CO2 incubator, with orbital shaking (170 rpm) when required. *Escherichia* coli NEB 5-α (New England Biolabs) and E. coli pir+ were employed for plasmid assembly and propagation. *E. coli* strains were grown aerobically in Lysogeny Broth (LB) at 37 with orbital shaking (220 rpm). Plasmids were introduced into *S. alvi* by electroporation, and into *Gilliamella* by conjugation, following previously described procedures^42^. When appropriate, antibiotics were added to culture media for selection and plasmid maintenance at the following concentrations: spectinomycin, 30 - 60 μg ml^-1^; ampicillin, 30 - 100 μg ml^-1^; and kanamycin, 25 - 50 μg ml^-1^, depending on whether the construct was maintained in *S. alvi/Gilliamella* or E. coli.

### Plasmid cloning

To expand the genetic toolbox for *S. alvi* and enable fine-tuned gene expression, we constructed a panel of plasmids carrying synthetic promoters of varying strengths sourced from a previously published study^51^. Promoters CP25, CP6, CP7, CP32, CP20, J23100, J23106, and J23116 were cloned into the pBTK570 backbone to generate plasmids pAC95, pAC82, pAC83, pAC84, pAC85, pAC86, pAC87 and pAC88, respectively. In each case, the original PA3 promoter driving *E2-crimson* in pBTK570 was replaced with the corresponding promoters using 5’ overhangs introduced by PCR primers. The pBTK570 backbone was amplified and linearized into two fragments using the primer pairs AC436/38 and AC435/39 for CP25; AC384/38 and 386/39 for CP6; AC388/38 and 387/39 for CP7; AC390/38 and 389/39 for CP32; AC392/38 and 391/39 for CP20; AC394/38 and 393/39 for J23100; AC396/38 and 395/39 for J23106 and AC398/38 and 397/39 for J23116.

To generate the arabinose-responsive strains, we built the plasmids pAC51b, pAC51c, pAC51g, pAC51i and pAC51j. Plasmid pAC51b was constructed by amplifying the *araC* gene from pBAD33 using primers AC271 and AC272, and inserting it into the pAC17v5b backbone, which was linearized into two fragments by PCR with primer pairs AC273/49 and AC274/50. Plasmid pAC51c was assembled by cloning a chemically synthesized pBAD25 promoter sequence, amplified with primers AC275 and AC276, into the pAC17v5a backbone, also linearized into two fragments using primer pairs AC019/059 and AC022/060. Plasmid pAC51g was generated by amplifying *araC* from pAC51b using primers AC507 and AC09, and inserting it into the intermediate plasmid pAC51e, which was linearized with primer pairs AC59/69 and AC504/60. Plasmid pAC51e itself was built by cloning the E2-crimson gene, amplified from pBTK570 with primers AC499 and AC501, into the pAC51c backbone, which was linearized with primer pairs AC503/059 and AC504/060. Plasmid pAC51i was assembled by rejoining the pAC51b backbone, amplified as three fragments with primer pairs AC516/010, AC18/050 and AC049/517. During amplification, the promoter CP25 was replaced by CP6 using the 5’ overhang of primer AC517. Plasmid pAC51j was constructed by inserting the *gfp* gene, amplified from pAC51c with primers AC518 and AC009, into the pAC51b backbone linearized with primer pairs AC049/069 and AC274/050.

To generate the standardized suicide plasmids for homologous recombination in *S. alvi*, we constructed plasmids pAC62, pAC98, pAC114 and pAC115. Plasmid pAC62 was built by amplifying the R6K origin of replication from pKD3 using primers AC351 and AC352 and inserting it into pBTK570, which was linearized with primers AC010 and AC350. Plasmid pAC98 was assembled by amplifying the *kanR* cassette from pBTK519 with primers AC010 and AC447 and cloning it into pAC62 linearized with primers AC018 and AC448. Plasmid pAC114 was generated by amplifying the *ampR* cassette from pAC08 using primers AC010 and AC541 and inserting it into pAC98 linearized with primers AC018 and AC540. Plasmid pAC115 was constructed by cloning the *tetR* gene, amplified from the genome of *S. alvi* using primers AC531 and AC532, into the pAC98 backbone linearized with primers AC533 and AC534, together with a short double-stranded DNA fragment formed by annealing complementary oligonucleotides AC538 and AC539.

Fragments were assembled using the NEBuilder HiFi DNA Assembly kit according to the manufacturer guidelines. Sequences of cloning primers and synthesized DNA can be found in **S1 Table** and **S2 Table**, respectively. Full plasmid maps of suicide backbones and arabinose constructs are depicted in S1 Fig and S2 Fig, respectively. The pBAD33 vector was kindly provided by Prof. David Tirrell (California Institute of Technology, United States)^65^. The pAC17v5a (Addgene plasmid No. 197413), pAC17v5b (No. 197414) and pAC08 (No. #197402) plasmids were previously built by our group^42^. The pBTK570 (No. 110615) and pBTK519 (No. 110603) plasmids were a gift from Prof. Jeffrey Barrick^40^. The pKD3 (No. 45604) plasmid was a gift from Barry L. Wanner^66^.

### Engineering of *S. alvi* for arabinose biosensing

To enable *S. alvi* to import arabinose, we integrated the *araE* transporter into its genome via homologous recombination^41^. To generate the *araE* insertion construct, we first retargeted the pAC98 backbone, carrying the *kanR* cassette, to the *tetC* locus. Approximately 1 kb-homology arms flanking *tetC* were amplified from the genome of *S. alvi* using primer pairs AC437/438 and AC439/440, and inserted into the backbone, which was linearized into two fragments using the primers AC010/216 and AC408/409 (**S1 Fig**). These standardized primers can be reused for retargeting any of the suicide plasmids, provided that the following constant tail sequences are appended to the primers used to amplify the homology arms: 5’-cacttaacggctgacatggg-3’ and 5’-catgaccaaaatcccttaacgtg-3’ for one arm, and 5’- cggatttacaattcgtcgtgc-3’ and 5’-catctgaatcatgcgcggat-3’ for the other.

Using the *tetC*-retargeted plasmid, we generated the final araE insertion fragment by overlap PCR. The *araE*coding sequence was amplified from the genome of *E. coli* using primers AC449/450, while the homology arms and *kanR* fragments were amplified from the retargeted plasmid with primer pairs AC446/477 and AC445/462. PCR products were combined at equimolar ratios (0.1 pmol each) in 20 μl of MiliQ water and assembled using 25 μl of high-fidelity DNA polymerase (Phanta master mix, Vazyme). The first overlap-extension stage was performed with the following conditions: 5 min at 95 °C, followed by 15 cycles of 20 sec at 95 °C, 1 min at 58 °C and 2.5 min at 72 °C; and a final extension for 5 min at 72 °C. The resulting assembly was resolved on an agarose gel, and the band of the expected size was gel purified. A second amplification stage was then performed by mixing 2 μL of the purified overlap product with 2 μl of each of the 10 μM primers AC445 and AC446, 20 μl of MiliQ water and 25 μl of high-fidelity polymerase. Amplification was carried out with the following conditions: 5 min at 95 °C, 30 cycles of 20 sec at 95 °C, 30 sec at 58 °C and 2.5 min at 72 °C; followed by a final extension for 5 min at 72 °C.

The final overlap product was purified and 1 μg of linear DNA was electroporated into S. alvi. Successful genomic integration of the araE gene was confirmed by Sanger sequencing of the tetC locus. The resulting strain was then transformed sequentially by electroporation with plasmid pAC51b followed by pAC51c, yielding the biosensor strain S. alvi ac02.

### Flow cytometry

Cell fluorescence was quantified using a NovoCyte flow cytometer (Agilent) and the NovoExpress software (version 1.4.1). To assess arabinose response of *S. alvi* ac02 and its derivative strains, as well as measuring gene expression levels of *S. alvi* carrying the synthetic low-strength promoters, single colonies were streaked as bacterial lawns on TSA plates supplemented with the appropriate antibiotics and incubated for 3-4 days as described above. Cells from each strain were then scraped from the agar surface, resuspended in liquid TSB to an optical density of 1.0 (OD600 = 1), and used to inoculate five biological replicates by diluting 1:300 in fresh TSB supplemented with varying arabinose concentrations when required. To assess the specificity of the response to arabinose, TSB was supplemented with 0.7 mM of either xylose, ribose, or arabinose. After 3 days of growth, cultures were diluted 1:10 in sterile PBS for flow cytometry analysis.

Fluorescent cells were identified using a two-step gating strategy: first, bacterial cells were selected on a FSC-H/SSC-H plot; single cells were then isolated on a FSC-A/FSC-H plot. Fluorescence of singlets was subsequently recorded using the FITC-H channel for GFP (ex. 488 nm – em. 530/30 nm) and the PE-Texas Red-H channel (ex. 561 nm – em. 615/20 nm) for E2- crimson. Mean fluorescence values were calculated from a minimum of 10,000 bacterial events per sample. An example of the gating strategy is provided in **S10 Fig**.

### Determination of the honeybee gut volume

To estimate honeybee gut volume, we quantified water loss following desiccation of dissected guts from 20 bees co-colonized with *S. alvi* ac02 and either *G. apicola* ESL309 or *G. apis* ESL169, collected across multiple cages from the experiment described below. At day 7 post-colonization, 10 bees were sampled from the treatment in which bees were fed sugar water (SW) supplemented with 210 μM arabinose, and 10 additional bees were sampled from the treatment in which bees were fed SW and pollen, to assess whether pollen consumption influenced gut volume.

Each dissected gut was transferred into a pre-weighted 1.5 ml Eppendorf tubes, and tubes were weighed again to determine fresh gut mass. Tubes were then left open and placed in an oven at 65 °C (fan 100%). After 22h, tubes were closed, cooled at room temperature for ∼30 min, reopened, and weighed to obtain the dry mass. Tubes were returned to the oven for an additional 4 h and weighed once more to ensure that the dry mass had stabilized. Gut volume in μl was estimated assuming a density of 1 mg μl^-1^ and calculated as the mass difference between fresh and dry gut weigh. As no significant difference in gut volume was observed between bees fed pollen and those not fed pollen, we used the overall mean volume of 19.1 μl for subsequent calculation.

### GC-MS quantification of arabinose

Arabinose levels in the gut were compared between MD bees and those colonized with a synthetic community of 25 isolates that encompass highly prevalent and abundant species within the gut microbiota of honeybees comprising the Bartonella, Bifidobacterium, *Bombilactobacillus*, *Commensalibacter*, *Frischella*, *Gilliamella*, Lactobacillus and *Snodgrassella* genera^3^. Bees were given sugar water and sterilized pollen ad *libitum* for ten days.

The experiment was duplicated using bees from different hives with 4 bees sampled per replicate per condition. Full hindguts were collected from anesthetized bees and homogenized with glass beads in 500 μl of deionized water. An aliquot of 125 μl was then centrifuged for 20 min at 4 °C and 20,000 g and the supernatant was stored at −80 °C.

Gut homogenates and calibration curves of individual sugars were prepared identically. A 5 μl mixture of D-norleucine and vanillic acid internal standards was spiked into 20 μl of each sample immediately before extraction with 100 μl of cold acetonitrile:methanol (1:1, v/v). Samples were stored at −20 °C for 2 h, then centrifuged at 18,000 g for 20 min at 4 °C. The supernatant was transferred to a new plate and dried in a speed vacuum concentrator. Samples were derivatized with 50 μl of 20 mg ml^-1^ methoxyamine hydrochloride in pyridine for 90 min at 34 °C, followed by 50 μl of N-Methyl-N-(trimethylsilyl)trifluoroacetamide (MSTFA) for 120 min at 45 °C.

Samples were then randomized and 1 μl was injected via autosampler into an Agilent 8890/5977B GC-MSD with a 2:1 split ratio and inlet temperature of 280 °C. Chromatography was performed using a VF-5MS column (30 m × 0.25 mm × 0.25 μm) with 1 ml min^-1^ helium flow rate. The oven was held for 2 min at 125 °C, increased at 3 °C/min to 150 °C, 5 °C/min to 225 °C, 15 °C/min to 300 °C, 25 °C/min to 310 °C and held for 3.3 min. The mass spectrometer was run in scan mode over a mass range of 50-550 Da at a scan rate of 3.2 scans/s. Sugar peak areas were quantified using Agilent MassHunter Quantitative Analysis software (version 10.0). Metabolite peak areas were normalized across samples using the median internal standard areas, and a linear regression curve of metabolite concentration vs peak area was calculated for each sugar.

### Plasmids maintenance measurement in vitro

For the assessment of pAC51b and pAC51c stability, single colonies of *S. alvi* ac02 were inoculated into liquid TSB and grown under three conditions, each with 10 biological replicates: (*i*) 30 μg ml^-1^ spectinomycin, 30 μg ml^-1^ ampicillin and 25 μg ml^-1^ kanamycin (+ ab); (*ii*) 25 μg ml^-1^ kanamycin only (- ab); or (*iii*) 25 μg ml^-1^ kanamycin and 0.07 mM arabinose (- ab + ara). After 3 days of incubation, cultures were serially diluted and plated onto both selective and non-selective TSA. Plasmid maintenance was quantified as the proportion of colonies that grew on selective plates relative to the total number of colonies on non-selective plates, corresponding to cells maintaining both plasmids.

### Honeybee rearing and gut colonization

Microbiota-deprived (MD) *Apis mellifera carnica* bees were obtained from outdoor colonies maintained at the University of Lausanne (VD, Switzerland), following previously described procedures^12^. Briefly, mature pupae were sampled from brood frames and transferred to sterilized plastic emergence boxes. Pupae were incubated at 35 with 75 % humidity for 3 days. Emerging adults were supplied with sterile 1:1 (w/v) sucrose solution.

To confirm the absence of microbial contaminants, hindguts of two newly emerged bees per box were dissected and homogenized in 1 ml sterile PBS. Homogenates were plated onto nutrient agar (NA), columbia blood agar (CBA), brain-heart infusion agar (BHI), LB and MRSA plates and incubated under aerobic (NA, LB), microaerophilic (CBA) and anaerobic (BHI, MRSA) conditions. Emergence boxes were excluded if any microbial growth was detected on the corresponding plates.

For mono-colonization, MD bees were individually fed 5 μl of *S. alvi* ac02 resuspended at OD600 of 0.1 in 1:1 (v/v) PBS:sucrose solution. For co-colonization with the native gut community, gut homogenate stocks were prepared by pooling equal volumes of homogenized hindguts from five hive bees. The colonization inoculum was generated by mixing S. alvi ac02 at OD600 of 0.1 with a 1:10 (v/v) dilution of the gut homogenate stock in PBS:sucrose solution. For co-colonization with *G. apicola* ESL309 and *G. apis* ESL169, bees were fed a mixture containing *S. alvi* ac02 and either of the *Gilliamella* species to a final OD600 of 0.1 and 1, respectively, in PBS:sucrose solution. *Gilliamella* strains carried the plasmid pAC09 (Addgene plasmid No. 197403)^42^ encoding the E2-crimson fluorescent protein to enable in vivo visualization.

Following colonization, bees were maintained in sterile cup cages at 32 and 75 % humidity. For fecal bacterial load assays, mono-colonized bees were supplied with sugar water containing 85 μM arabinose, and either 30 μg ml^-1^ ampicillin plus 30 μg ml^-1^ spectinomycin (‘+ ab’) or no antibiotic (‘- ab’, ‘- ab + gc’). To assess the *in vivo* biosensor response to arabinose, mono-colonized bees received sugar water supplemented with 30 μg ml^-1^ of each ampicillin and spectinomycin, together with 0, 10, 100 or 1,000 μM arabinose. Bees co-colonized with *S. alvi* ac02 and *Gilliamella* were given one of three diets: (*i*) sugar water with 60 μg ml^-1^ ampicillin, (*ii*) sugar water with 60 μg ml^-1^ ampicillin and 210 μM arabinose, or (*iii*) sugar water with 60 μg ml^-1^ ampicillin and pollen provided ad *libitum*.

As pollen exhibits autofluorescence primarily in the red spectrum, which may interfere with E2-crimson signal detection, pollen was removed from cages two days prior to gut dissection. All supplemented feeding solutions (*i.e*. antibiotics and arabinose) were replaced with freshly prepared solutions every 3 days to maintain consistent molecule concentrations.

### Bacterial load quantification via fecal sampling

To quantify the fecal abundance of *S. alvi* ac02, fecal material was collected from bees at 7 and 14-day post colonization with the engineered cells. Bees housed in cup cages were stunned with CO2 and subsequently placed on ice to immobilize them. While anesthetized, gentle pressure was applied manually along the abdomen from the anterior end toward the stinger to induce defecation directly into the cap of sterile 1.5 ml Eppendorf tubes. Samples were maintained on ice throughout the collection period.

Fecal matter that was sufficiently liquid was immediately serially diluted 1:10 (v/v) in sterile PBS. For more viscous samples, feces were first resuspended in 4 μl of sterile PBS before performing the 1:10 serial dilution. Diluted samples were plated on TSA agar supplemented with 30 μg ml^-1^ spectinomycin, 30 μg ml^-1^ ampicillin and 25 μg ml^-1^ kanamycin to selectively recover *S. alvi* ac02. Colony forming unit (CFUs) were counted after 3-4 days of incubation.

For the assessment of pAC51b and pAC51c stability in vivo, resuspended feces were also serially diluted and plated onto both selective (ampicillin or spectinomycin) and non-selective TSA. Plasmid maintenance levels of each plasmid was quantified as the proportion of colonies that grew on the corresponding selective plates relative to the total number of colonies on non- selective plates.

### Gut microscopy analysis

To prepare samples for microscopy, honeybee hindguts were collected seven days after inoculation with engineered *S. alvi* and *Gilliamella,* a time point selected to ensure sufficient biosensor abundance for reliable sugar detection, and immediately immersed in 4 % paraformaldehyde in PBS. Tissues were fixed overnight at 4 with gentle rotation, followed by three 30-min washes at room temperature in PBS. Samples were then permeabilized and stained overnight at 4 in the dark with 5 μg ml^-1^ 4,6-diamidino-2-phenylindole (DAPI) prepared in PBS containing 1% Triton X-100. A single master mix of sufficient volume for all samples was prepared and used for the entire staining procedure, ensuring that all samples were stained using the same DAPI solution and eliminating potential variation between staining batches. After staining, excess dye was removed by washing in PBS, and the ileum was dissected, mounted in PBS and overlaid with a coverslip (number #1.5).

Image acquisition was performed using a Nikon AX R laser scanning confocal microscope (Nikon, Japan) and the software Nikon NIS-Elements C (version 5.42.06). Images were acquired with a 60X oil objective at 2048x2048 pixel format size (field of view: 294.6278x294.6278 μm) and an averaging of 4 images. Pinhole size was set to 0.3 airy units, yielding an optical sectioning of 0.32 μm. Fluorescence acquisition was performed sequentially in two passes to minimize crosstalk:E2-crimson (ex. 561 nm–em. 623-750 nm) and GFP (ex. 488 nm–em. 499-551 nm), followed by DAPI (ex. 405 nm–em. 420-551 nm).

For quantitative fluorescence analysis, all ileums were imaged using identical laser intensity and detector gain settings. Gut regions with visible pollen grains were excluded from imaging. For each gut, three regions were imaged: anterior, mid-ileum, and posterior. For each region, Z-stacks were also collected over a maximum depth of 40 μm with 1 μm step increment. Fluorescence quantification was automated using a custom Fiji macro developed for this study (see **S1 Text**). The macro was executed in Fiji (ImageJ2, version 2.16.0). Briefly, bacterial areas were segmented by applying a GFP fluorescence threshold identical across images. The resulting selected areas were copied to the corresponding DAPI images. Within each GFP-segmented region, mean fluorescence intensity per unit area was then extracted for both GFP and DAPI channels. For host cell nuclei, regions of interest were instead segmented by applying an identical DAPI fluorescence threshold across images. The threshold was selected to preferentially identify the high-intensity DAPI signal from host cell nuclei while excluding the majority of bacterial DAPI signal. Mean DAPI fluorescence intensity per unit area was then quantified within each DAPI-segmented region. As DAPI fluorescence was reduced to a similar extent in both bacterial biofilm and host cell nuclei in pollen-fed bees, despite identical staining conditions, a DAPI normalization factor (*k* = 0.6) was calculated as the ratio of mean DAPI fluorescence in pollen-fed bees to that in the other diet conditions (**S11 Fig**). DAPI values from pollen-fed bees were then divided by k to obtain corrected DAPI values. The observed reduction in DAPI signal in pollen-fed bees may reflect effects of pollen-derived DNA or other pollen-associated material on DAPI availability or staining efficiency. GFP intensities were then normalized to the corrected DAPI intensities measured from the same segmented area to account for variation in cell density^67,68^. When appropriate, normalized values from the three ileum regions were averaged to obtain a single fluorescence measurement per bee gut.

## Acknowledgements

We acknowledge Lucy Genier for her support in the laboratory with molecular biology work. We thank Ly Pheak Chhun and Gonçalo Matos for helpful scientific discussions throughout the project. We also thank Georgia Petsiou and Laure Guerra for assistance with the bee experiments and Estelle Pignon for providing the *Gilliamella* strains bearing the pAC09 plasmid. We are grateful to Silvia Moriano Gutierrez for proofreading the manuscript, and to Méline Garcia for identifying the *Gilliamella* species of interest.

This work was supported by the NCCR Microbiomes (National Centre of Competence in Research; https://nccr-microbiomes.ch/), funded by the Swiss National Science Foundation (grant no. 225148; https://www.snf.ch/) to P.E and Y.S. as well as by the University of Lausanne (https://www.unil.ch/). A.I.P-L. was funded by an EMBO Postdoctoral Fellowship (ALTF 975- 2025; https://www.embo.org/). The funders had no role in study design, data collection and analysis, decision to publish, or preparation of the manuscript.

## Author contributions

A.C., P.E. and Y.S. conceived the study. A.C. designed the experiments and performed molecular biology work, confocal microscopy, image analysis for fluorescence quantification and bee experiments with support from F.Z. Suicide plasmids were designed and built by A.C. and T.S. Bee and *in vitro* experiments for the revision were carried out by A.I.P-L. GC-MS arabinose measurements and related bee experiments were carried out by A.Q. and T.H.G.N. Y.S. and P.E. supervised the project. A.C. wrote the manuscript with input from co-authors.

## Data availability

The plasmids developed in this study have been deposited in Addgene and are available *via* the following accession numbers: pAC62 (#249659), pAC98 (#249660), pAC114 (#249661), pAC115 (#249662), pAC51b (#249663) and pAC51c (#249664). Confocal microscopy images have been deposited in the BioImage Archive under the accession number S-BIAD2461. The FIJI macro for automated fluorescence quantification of bacterial biofilm can be found in the S1 Text file, as well as on Zenodo (https://doi.org/10.5281/zenodo.21925112). The FCS files underlying the flow cytometry data shown in Fig. 2a and S5 Fig. can be found on Zenodo (https://doi.org/10.5281/zenodo.22029676). Source data are provided with this manuscript in the S1 Data file.

## Supporting information

**S1 Data. Source data underlying plots shown in the study.**

**S1 Fig.**
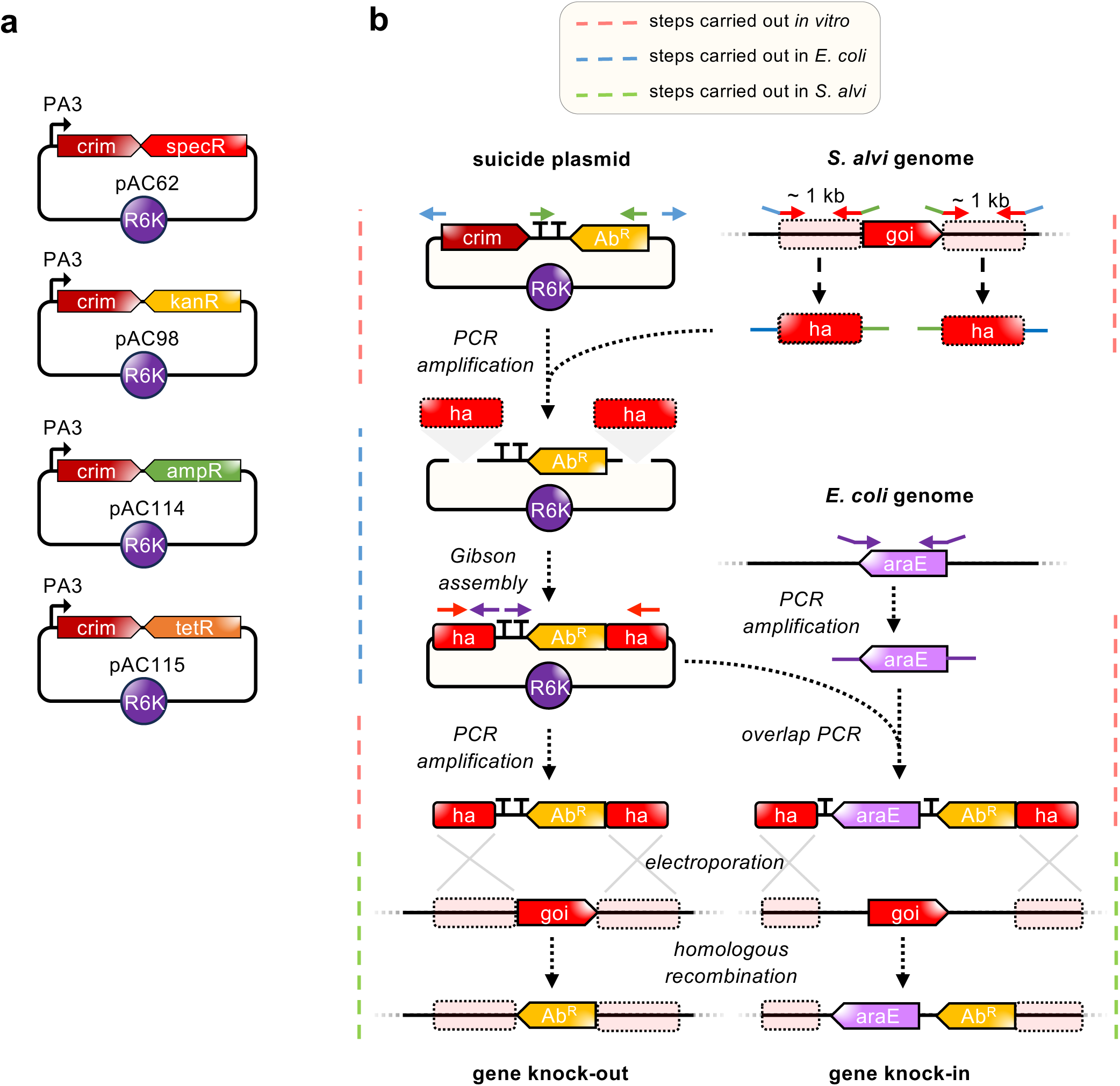
Genome engineering workflow for *S. alvi*. **(a)** Collection of standardized suicide plasmids developed in this study for easy and versatile genome engineering of *S. alvi*. The fluorescent marker E2- crimson (crim) carried by the suicide plasmids is removed following cloning of the homology arms, allowing easy confirmation of successful assembly as cells transformed with the plasmid containing homology arms will grow as non-fluorescent colonies. **(b)** Schematic of the genome engineering pipeline utilized for both gene knock-out of gene of interest (goi) and knock-in. Cloning of the homology arms flanking the insertion site of interest in the suicide plasmids can easily be performed by Gibson Assembly. For this, the standardized primer pairs AC_10/AC_216 (blue primers) and AC_408/AC_409 (green primers) allow efficient linearization of the suicide plasmids. Primers used for amplification of the homology arms (ha) must have standardized overhang sequences allowing assembly of the fragments with the linearized suicide plasmids. Using the resulting targeted plasmid, external primers of each homology arm can then be used to amplify the final insertion fragment to transform in *S. alvi* for homologous recombination-based genome engineering. The suicide plasmids bearing the R6K origin of replication can only be maintained in a *pir+* strain, ensuring that the acquisition of antibiotic resistance in the mutant strain results from the integration of the insert fragment rather than by replication of carry-over plasmid originating from the initial PCR reaction. For knock-in of genes, overlap PCR can be used to generate an insertion fragment from the retargeted plasmid, with another set of primers (purple primers).

**S2 Fig.**
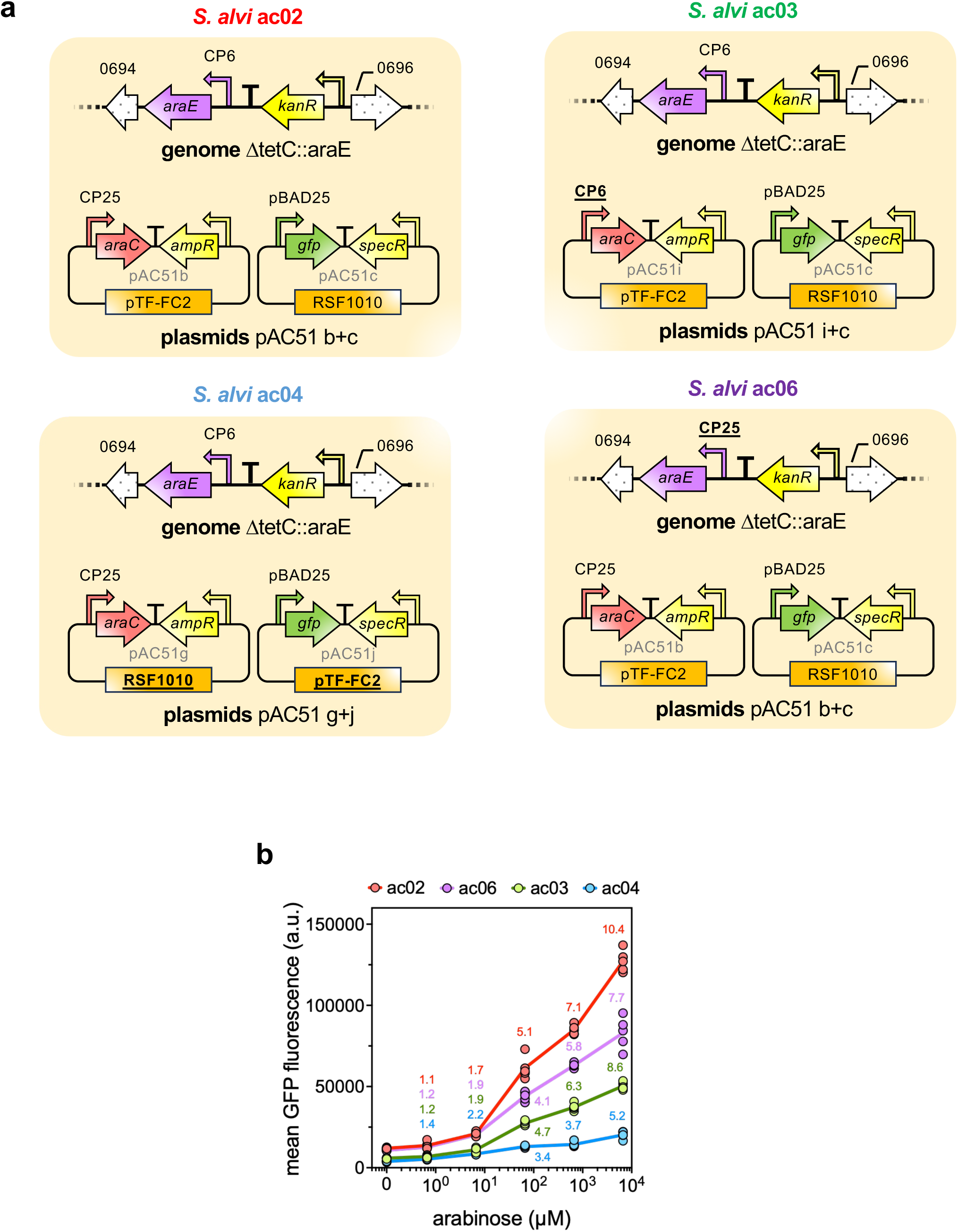
Variant *S. alvi* strains engineered for arabinose sensing display distinct response profiles. (**a**) Overview of genetic modifications of arabinose biosensor *S. alvi* strains. Differences in circuit design relative to the strain *S. alvi* ac02 are underlined and in bold. (**b**) The graph shows mean GFP fluorescence values from 5 biological replicates. Each replicate value was obtained from the average fluorescence of at least 9,000 bacteria measured by flow cytometry. The data lines are color-coded by strain, with the corresponding strain names (ac02-ac06) shown above the plot. Mean fluorescence fold-change relative to the uninduced condition is indicated in the same color. The data underlying this Figure can be found in the S1 Data file, sheets “S2b Fig”.

**S3 Fig.**
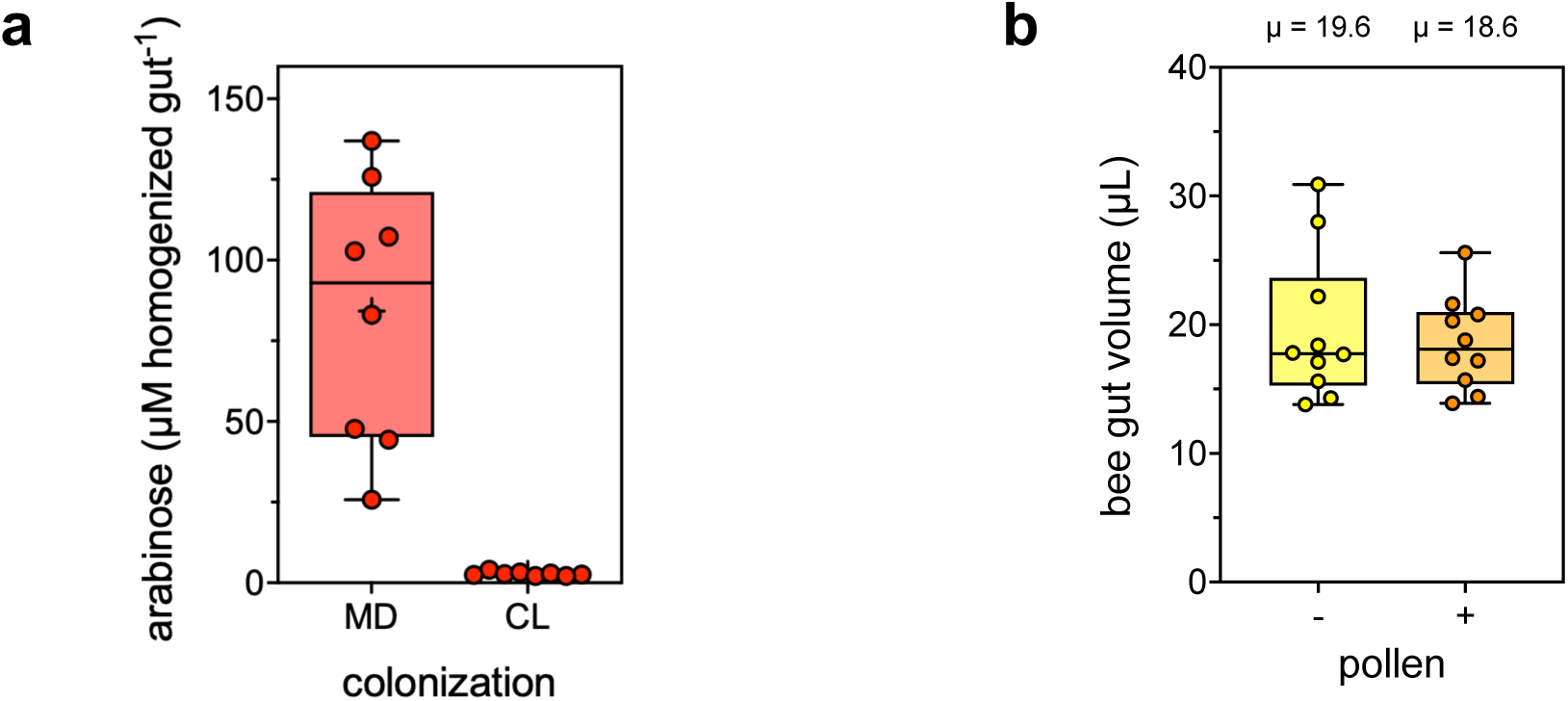
Quantification of arabinose concentrations in the honeybee gut. **(a)** Pollen-derived arabinose concentrations in bee guts. Box plots show median arabinose concentration measured by GC-MS of 8 bee guts homogenized in 500 µl of PBS. Bees were fed with sugar water and pollen. They were either microbiota-deprived (MD) or colonized (CL) for 7 days prior to dissection and arabinose quantification. **(b)** Determination of the honeybee gut volume. Box plots show median gut volumes calculated from the mass of water loss after desiccation of 10 guts collected from caged honeybees 7 days post-emergence. Bees were fed with (+) or without (−) pollen and sugar water *ad libitum*. The data underlying this Figure can be found in the S1 Data file, sheets “S3a Fig” and “S3b Fig”.

**S4 Fig.**
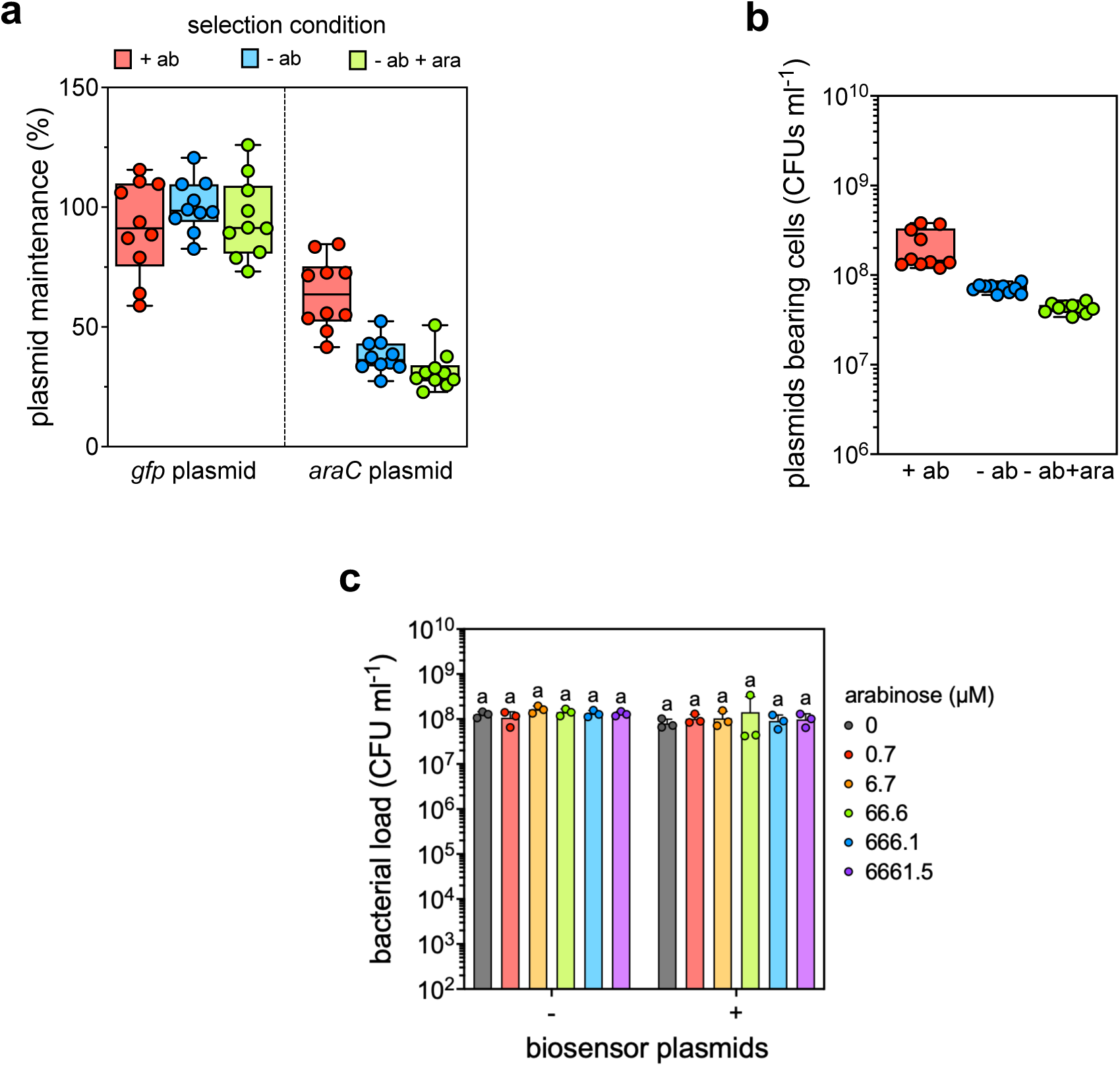
Stability of plasmids and growth characteristics of *S. alvi* ac02 *in vitro*. Maintenance and plasmid-bearing cell number of arabinose-inducible plasmids pAC51b and pAC51c in *S. alvi* ac02 in the absence of antibiotic selection. Box plots show median plasmid maintenance **(a)** and median concentration of plasmids-bearing cells **(b)** of 10 biological replicates. Each replicate represents a 3-day liquid culture grown in TSB with ampicillin and spectinomycin (+ab), TSA alone (-ab) or TSA with arabinose (-ab+ara). **(a)** Maintenance levels for pAC51b (*araC* plasmid) and pAC51c (*gfp* plasmid) were calculated as the fraction of CFUs growing on TSA with ampicillin or spectinomycin, respectively, relative to total CFUs on non-selective TSA. Incomplete maintenance of pAC51b in the +ab condition likely reflects the use of ampicillin selection, as β-lactamase-mediated degradation of ampicillin can reduce selection pressure during prolonged liquid culture and permit the expansion of plasmid-free cells. **(b)** Number of functional biosensor cells bearing the two plasmids was estimated by isolating CFUs growing on TSA with both antibiotics. **(c)** Growth assessment of *S. alvi* ac02 carrying the dual-plasmid system (pAC51b/c; +) compared to plasmid-free cells (−) following growth to stationary phase in the presence of different arabinose concentrations. Bar plots show mean of CFU counts ± standard deviation of 3 biological replicates. No significant differences in CFU counts were observed between plasmid or arabinose conditions (two-way ANOVA: effect of plasmids, F(1, 24) = 2.68, p = 0.114; effect of arabinose, F(5, 24) = 0.48, p = 0.785). The data underlying this Figure can be found in the S1 Data file, sheets “S4a Fig”, “S4b Fig” and “S4c Fig”.

**S5 Fig.**
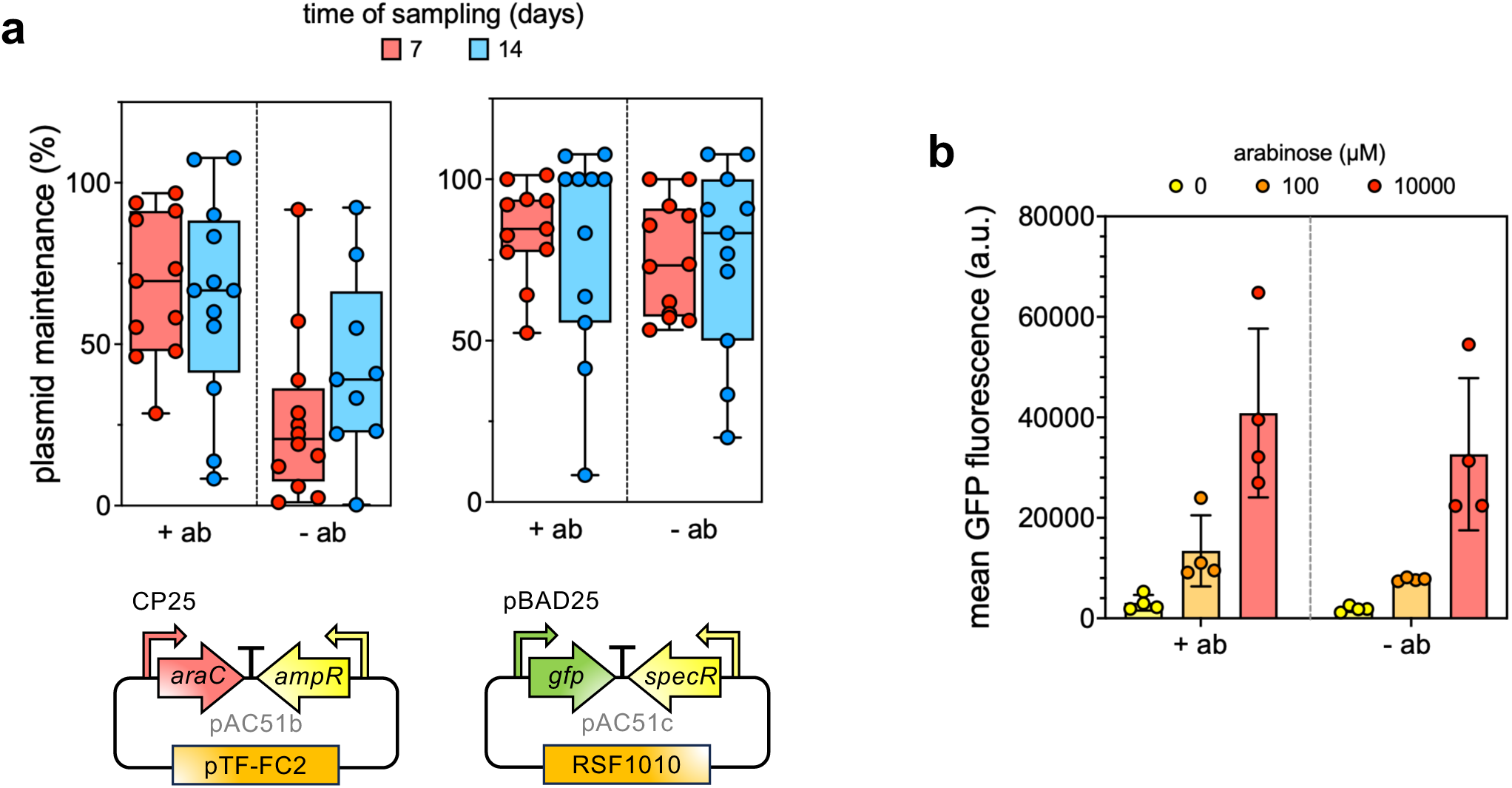
Stability of *S. alvi* ac02 *in vivo*. **(a)** Maintenance of arabinose-inducible plasmids pAC51b (left) and pAC51c (right) in *S. alvi* ac02 cells isolated from bee feces collected 7 (red) and 14 (blue) days after colonization. Bees were mono-colonized with the biosensor strain and fed sugar water supplemented with (+ab) or without (−ab) antibiotic. Box plots show median plasmid maintenance of at least 9 biological replicates. **(b)** The *S. alvi* ac02 strain remains functional after gut colonization. Colonies were randomly selected from the day 7 CFU-counting experiment after isolation on media supplemented with ampicillin and spectinomycin, ensuring retention of both plasmids. Cells originated from bees fed sugar water with (+ab) or without (−ab) antibiotic supplementation, and were grown in fresh liquid media with different concentrations of arabinose. Bar plots show mean GFP fluorescence values ± standard deviation from 4 biological replicates. Each replicate value was obtained from the average fluorescence of at least 9,000 bacteria measured by flow cytometry. The data underlying this Figure can be found in the S1 Data file, sheets “S5a Fig” and “S5b Fig”.

**S6 Fig.**
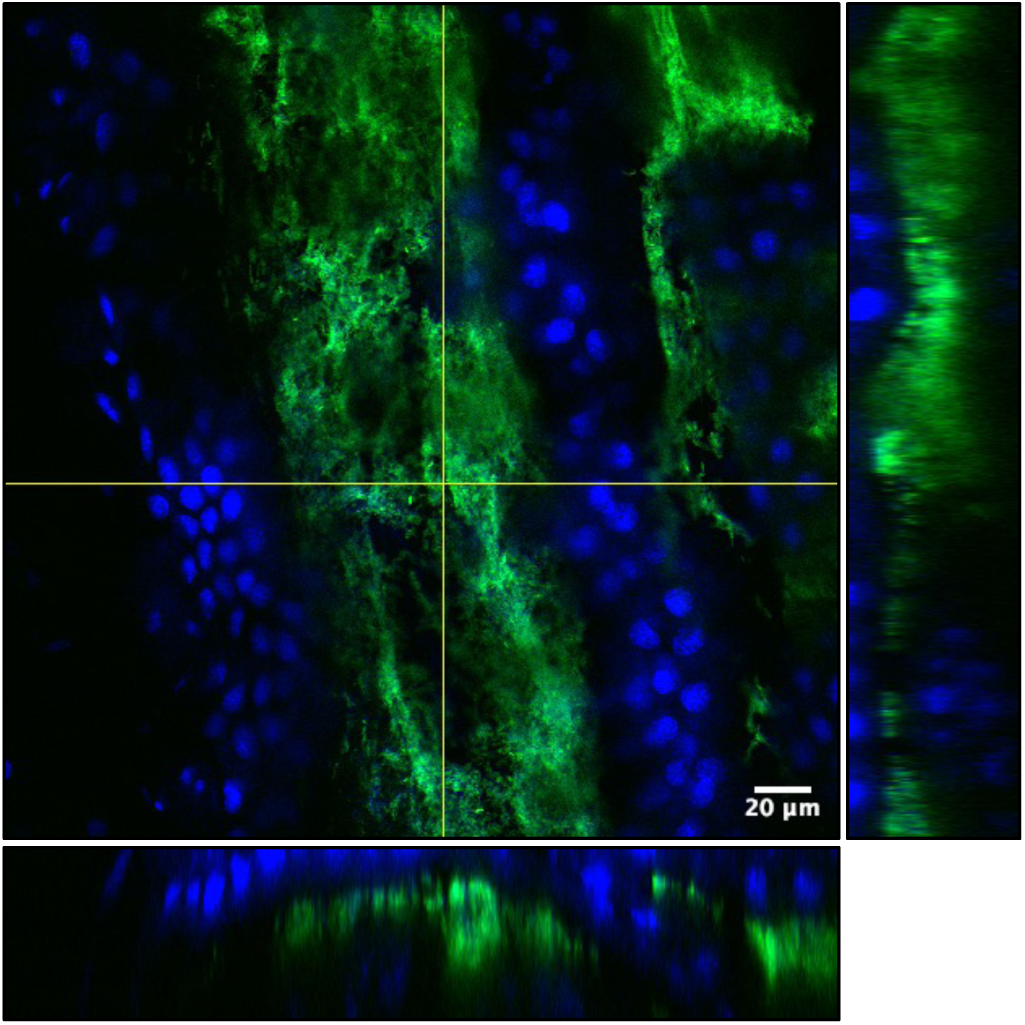
Example of a bacterial biofilm imaged from the epithelial attachment site to the honeybee gut lumen. Confocal microscopy image of a honeybee ileum colonized by *S. alvi* ac02. Blue and green channels are overlaid. The blue channel depicts DAPI staining of host and bacterial DNA. The green channel depicts GFP fluorescence from the engineered *S. alvi* ac02. Side panels show XY and YZ projections of 30 slices. Scale bar represents 20 µm.

**S7 Fig.**
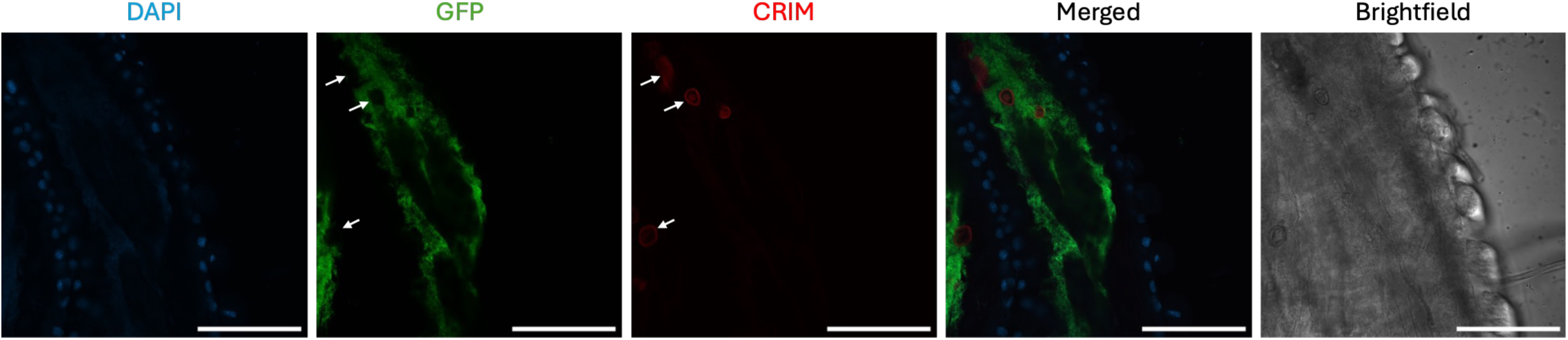
Pollen grains autofluorescence in the red spectrum. A confocal microscopy image of a honeybee gut colonized by engineered *S. alvi* ac02 in the ileum region. The bee was fed sugar water and pollen *ad libitum*. The blue channel depicts DAPI staining of host and bacterial DNA and the green channel shows GFP fluorescence from the engineered strain. Scale bar represents 100 µm. Localization of pollen grains are indicated by white arrows. Such images with visible pollen grains were not used to collect data shown in our manuscript but were taken as examples during test experiments carried out to optimize our protocols. Gut regions with visible pollen grains were otherwise excluded from imaging.

**S8 Fig.**
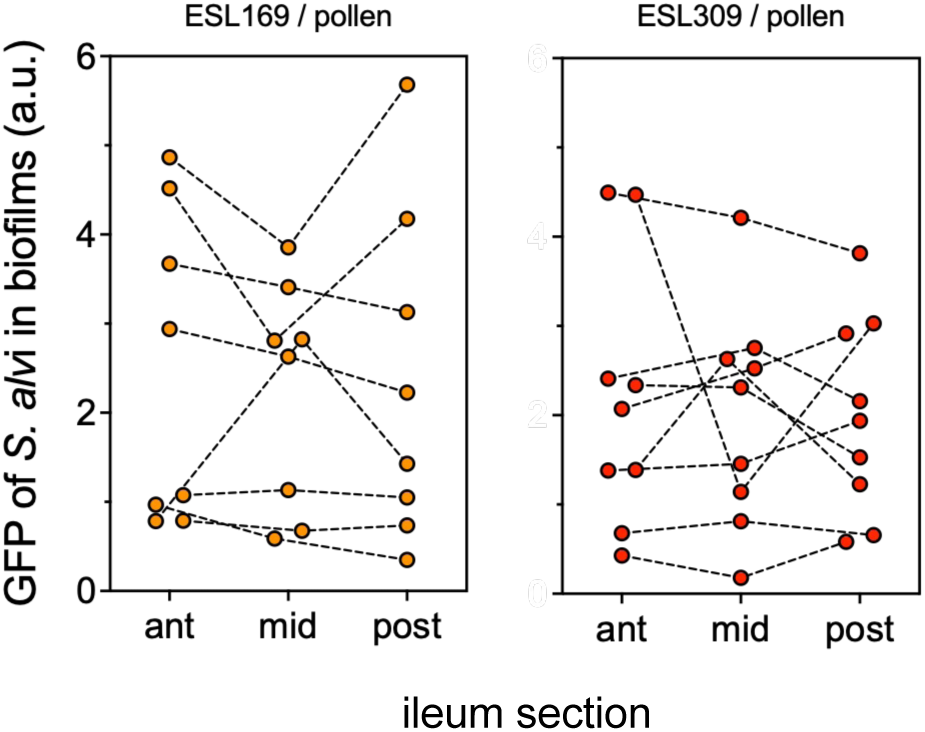
Concentration of pollen-derived arabinose is longitudinally uniform within each honeybee but highly heterogenous across individuals. Dot plot shows median value of GFP fluorescence quantified from *S. alvi* biofilms imaged in the gut of bees fed sugar water and pollen *ad libitum*. Each fluorescence value corresponds to the mean intensity averaged across the depth of the bacterial biofilm for each gut section. Paired samples are indicated with a dotted line. The data underlying this Figure can be found in the S1 Data file, sheet “S8 Fig”.

**S9 Fig.**
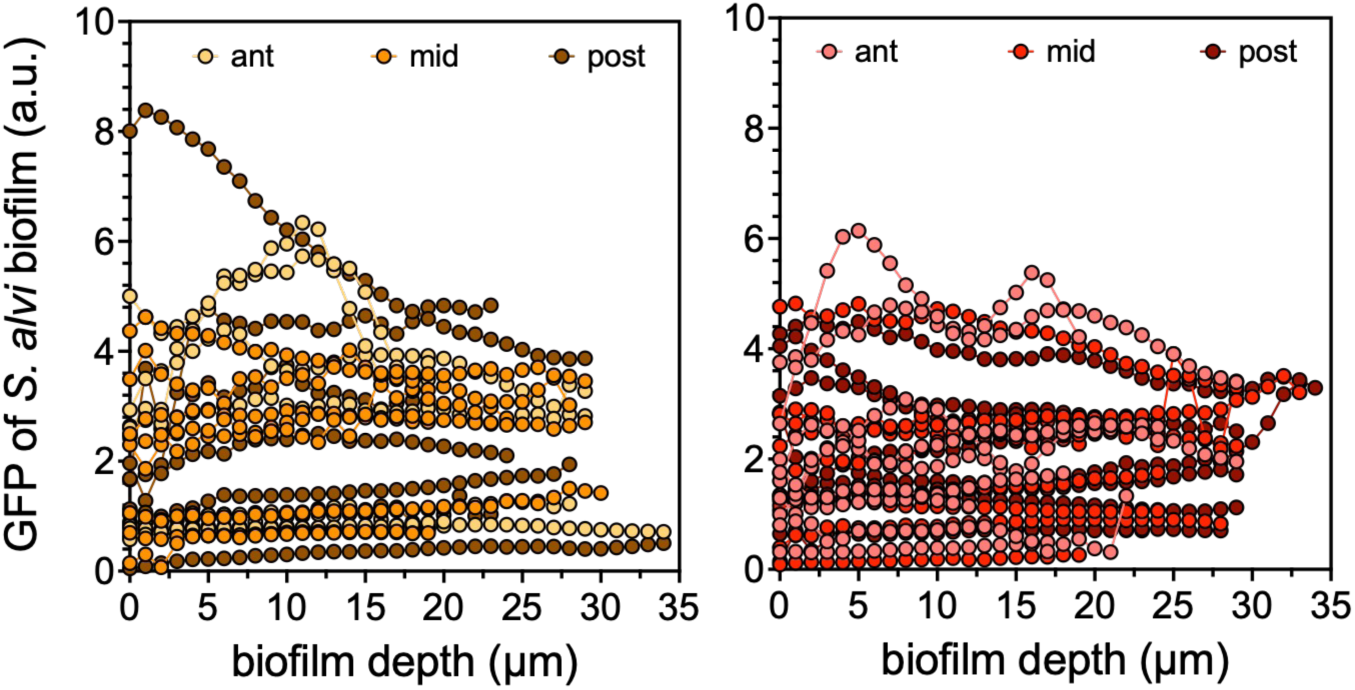
Arabinose derived from pollen breakdown distributes with high heterogeneity along a radial gradient across the depth of *S. alvi* ac02 biofilm when co-colonized with *G. apis* ESL169 (left) or *G. apicola* ESL309 (right). Graph shows mean GFP fluorescence of the biosensor according to the biofilm depth. For each bee, the graph displays the average signal for distinct ileum regions (anterior, mid-ileum, posterior). The data underlying this Figure can be found in the S1 Data file, sheet “S9 Fig”.

**S10 Fig.**
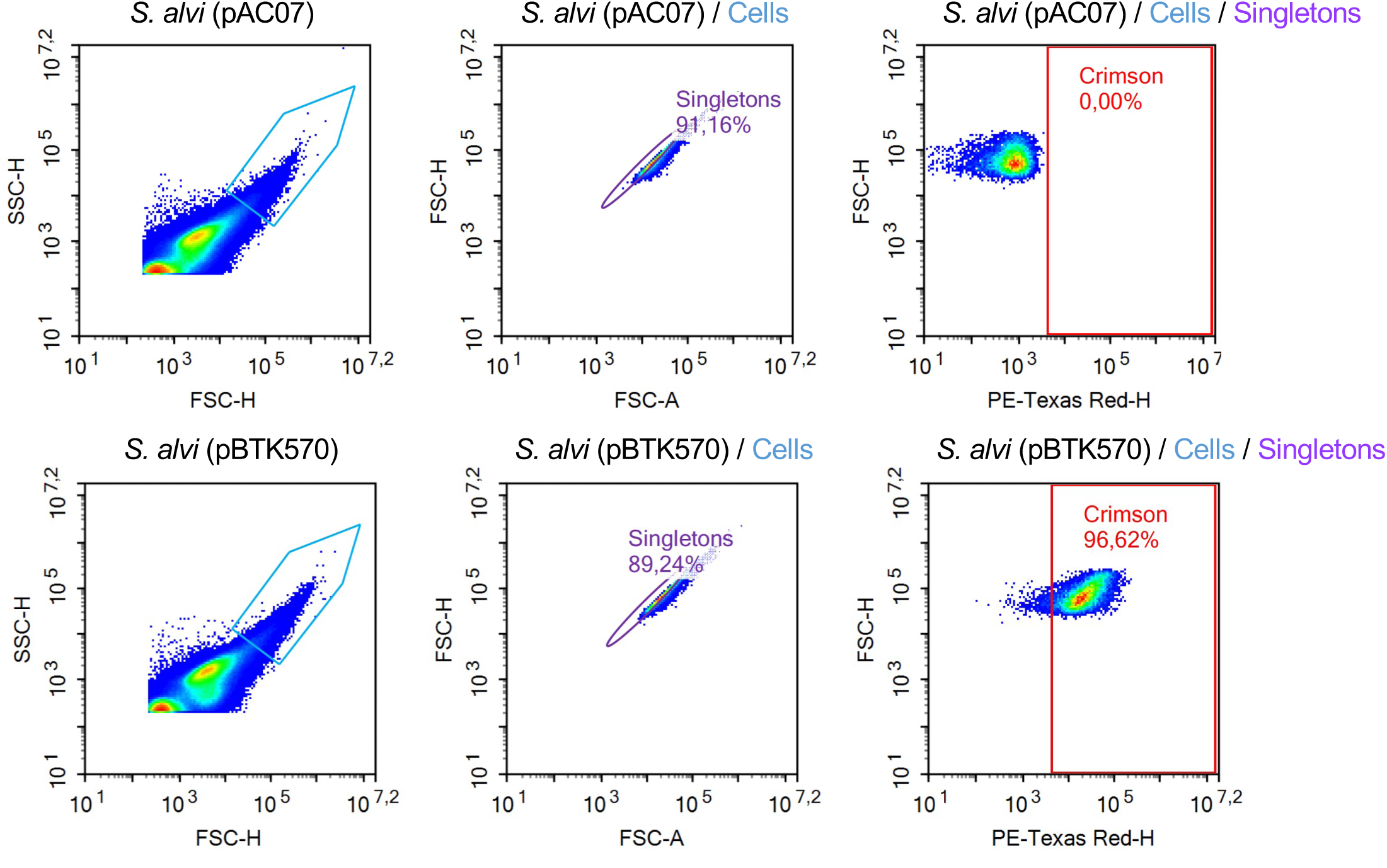
Cell gating for flow cytometry analysis. Fluorescent cells were identified using a two-step gating strategy: first, bacterial cells were selected on a FSC-H/SSC-H plot; single cells were then isolated on a FSC-A/FSC-H plot. Fluorescence of singlets was subsequently recorded using the FITC-H channel for GFP (ex. 488 nm – em. 530/30 nm) and the PE-Texas Red-H channel (ex. 561 nm – em. 615/20 nm) for E2-crimson. Mean fluorescence values were calculated from a minimum of 10,000 bacterial events per sample. Two examples of *S. alvi* cell gating are shown, with graphs representing pseudocolor plots. Quadrant limits were determined based on the measured fluorescence of reference cells bearing the empty backbone pAC07 (no fluorescence; top panel), or pBTK570 (E2-crimson alone; bottom panel).

**S11 Fig.**
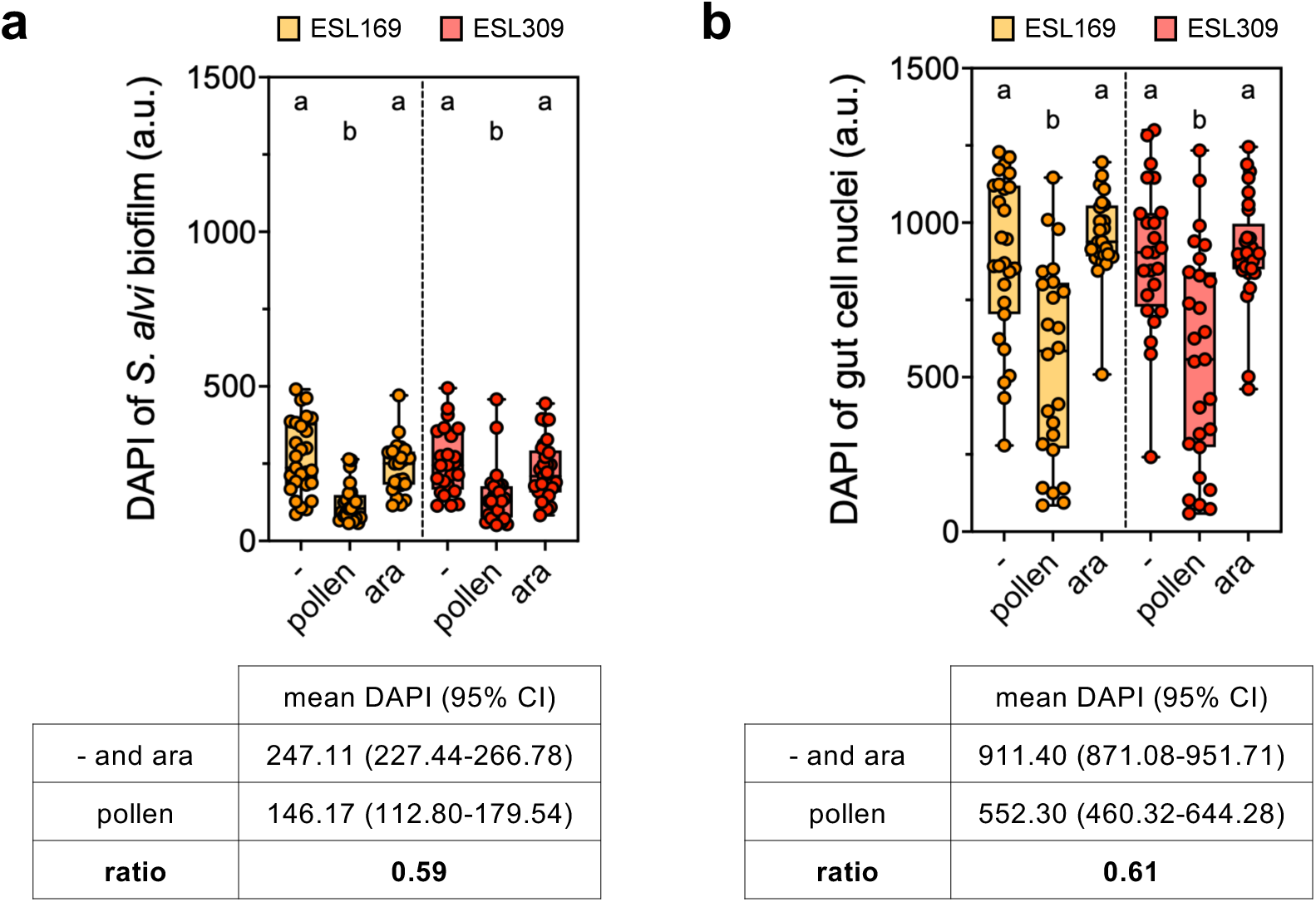
Pollen influences DAPI fluorescence in *S. alvi* biofilm and honeybee gut cell nuclei. Box plot shows median DAPI fluorescence of stained **(a)** *S. alvi* biofilms or **(b)** gut cell nuclei, imaged from honeybees fed sugar water (−), sugar water and pollen *ad libitum* (pollen) or sugar water supplemented with 210 µM arabinose (ara). Individuals were co-colonized by *S. alvi* ac02 and *G. apis* ESL169 (orange) or *G. apicola* ESL309 (red). For **(a)**, bacterial biofilm regions were identified by applying an identical GFP fluorescence threshold across images, and DAPI fluorescence was quantified within the resulting GFP- segmented regions. For **(b)**, host cell nuclei were identified by applying an identical DAPI fluorescence threshold across images; the threshold was selected to preferentially identify the high-intensity DAPI signal of host nuclei while excluding the majority of bacterial DAPI signal. DAPI fluorescence was then quantified within the obtained regions. Each fluorescence value corresponds to the overall mean intensity of a gut section (anterior, mid-ileum, or posterior). All samples were stained simultaneously with a single batch of staining solution (5 µg ml^-1^ DAPI in PBS with 1% Triton X-100), which was prepared once in sufficient volume for all samples. Different letters indicate significant differences in fluorescence intensities across colonization and diet conditions, with adjusted *p*-value < 0.05 (Kruskal-Wallis test). Mean DAPI fluorescence for the indicated conditions and the DAPI ratio are indicated below. The DAPI ratio was calculated as the mean DAPI fluorescence in pollen-fed bees divided by the mean DAPI fluorescence in the other diet conditions (- and ara). The data underlying this Figure can be found in the S1 Data file, sheet “S11 Fig”.

**S1 Table.**
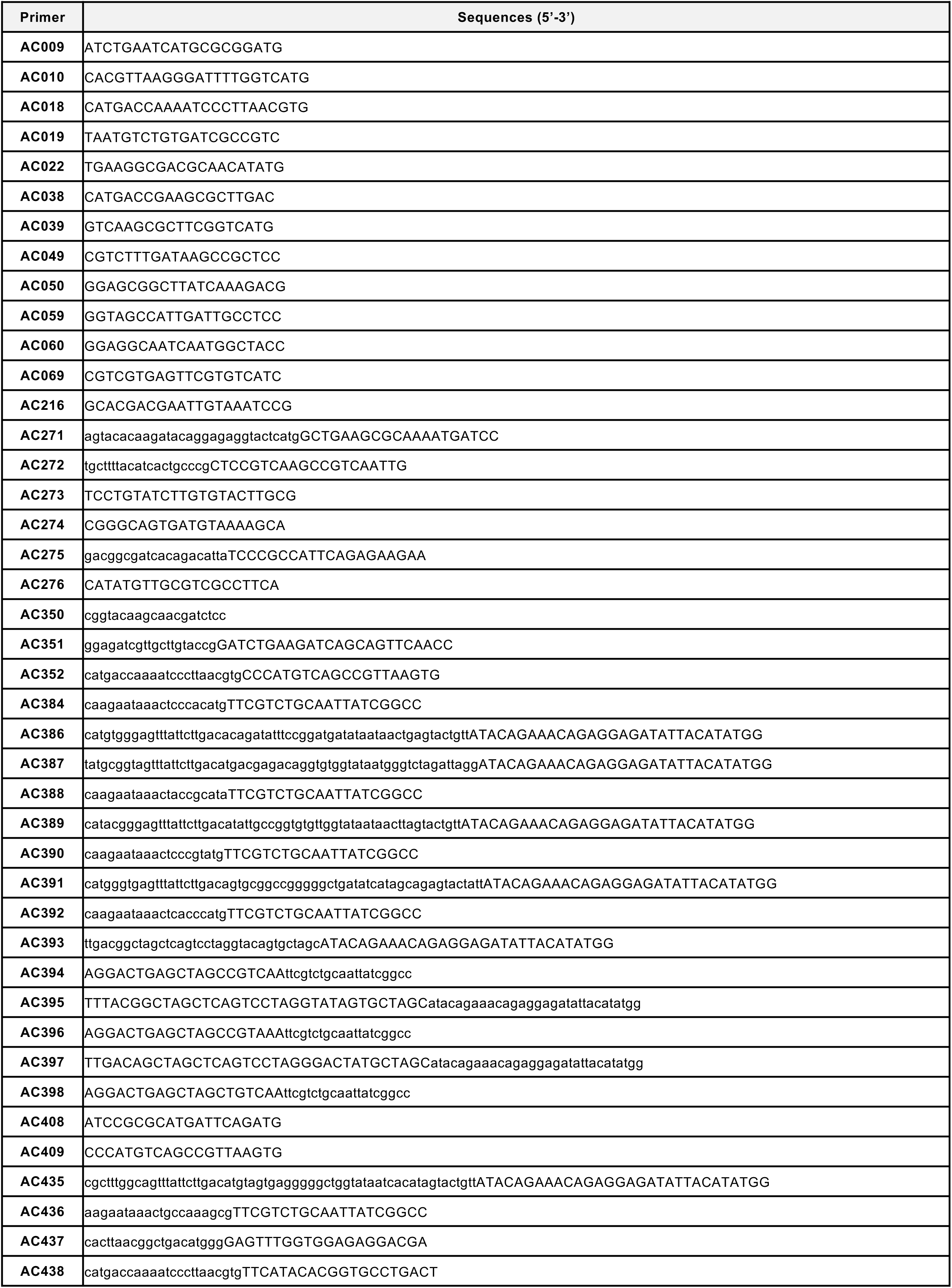
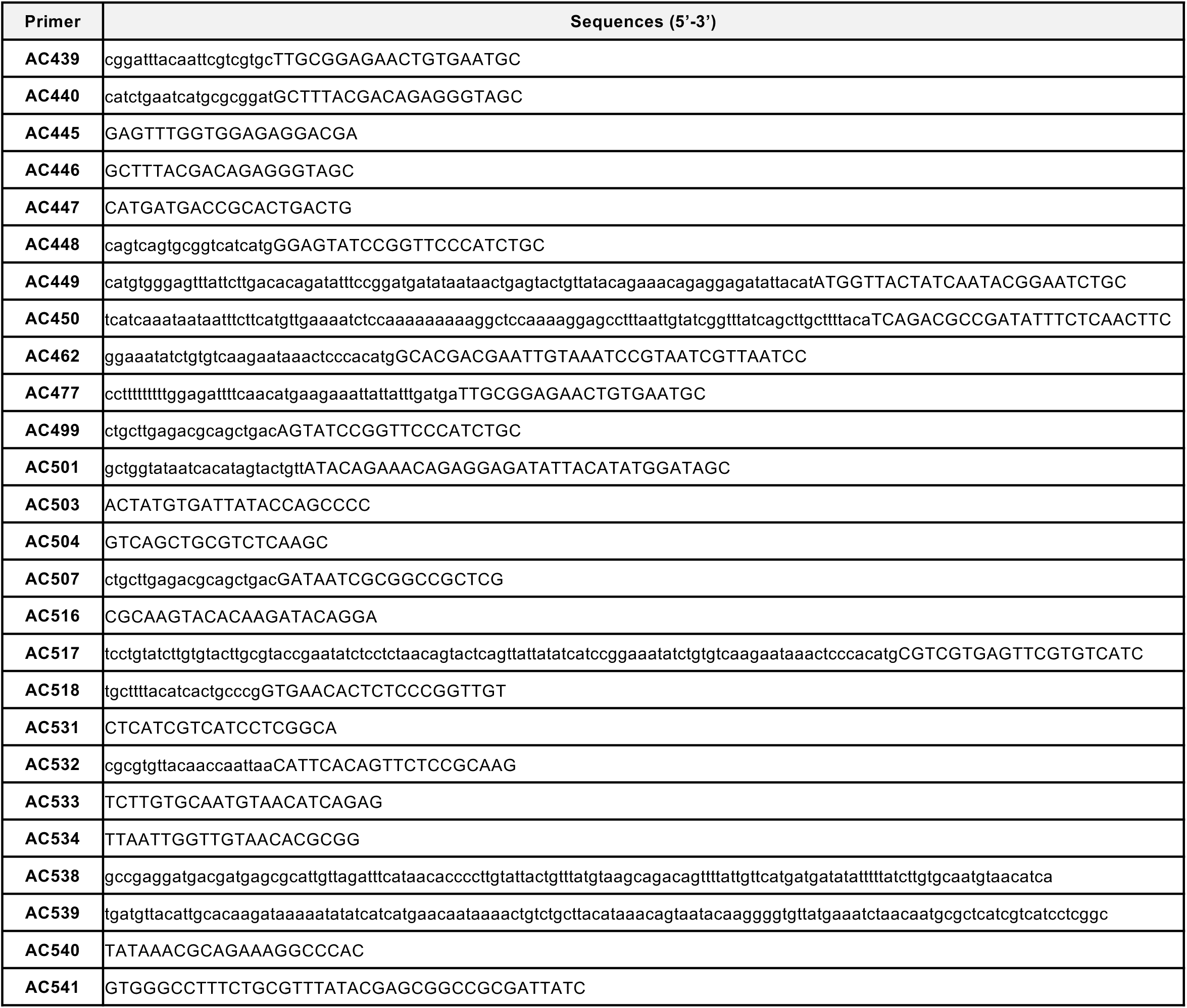
Primer sequences used in this study. Uppercase letters denote priming sequences, while lowercase letters represent Gibson assembly overhangs providing homology regions.

**S2 Table.**
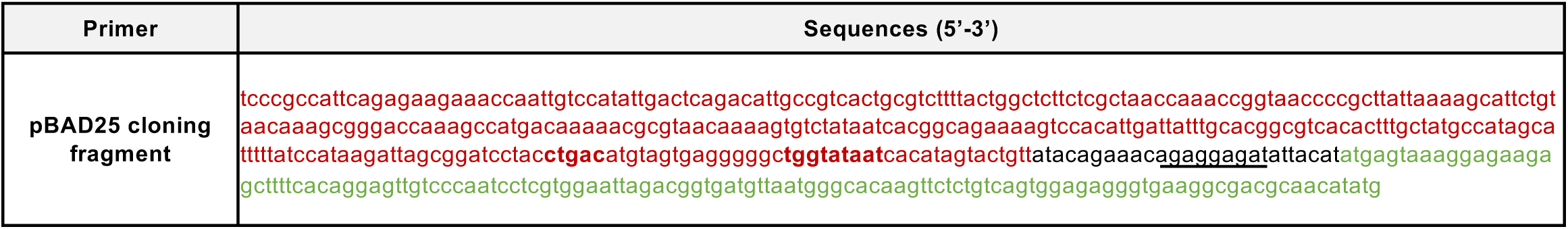
Synthesized DNA sequence used in this study. The pBAD25 promoter sequence is shown in red, the GFP fragment in green, the ribosome-binding site (RBS) is underlined, and the −35 and −10 promoter elements are indicated in bold.

**S1 Text. Fiji macro for automated fluorescence quantification of bacterial biofilm.**

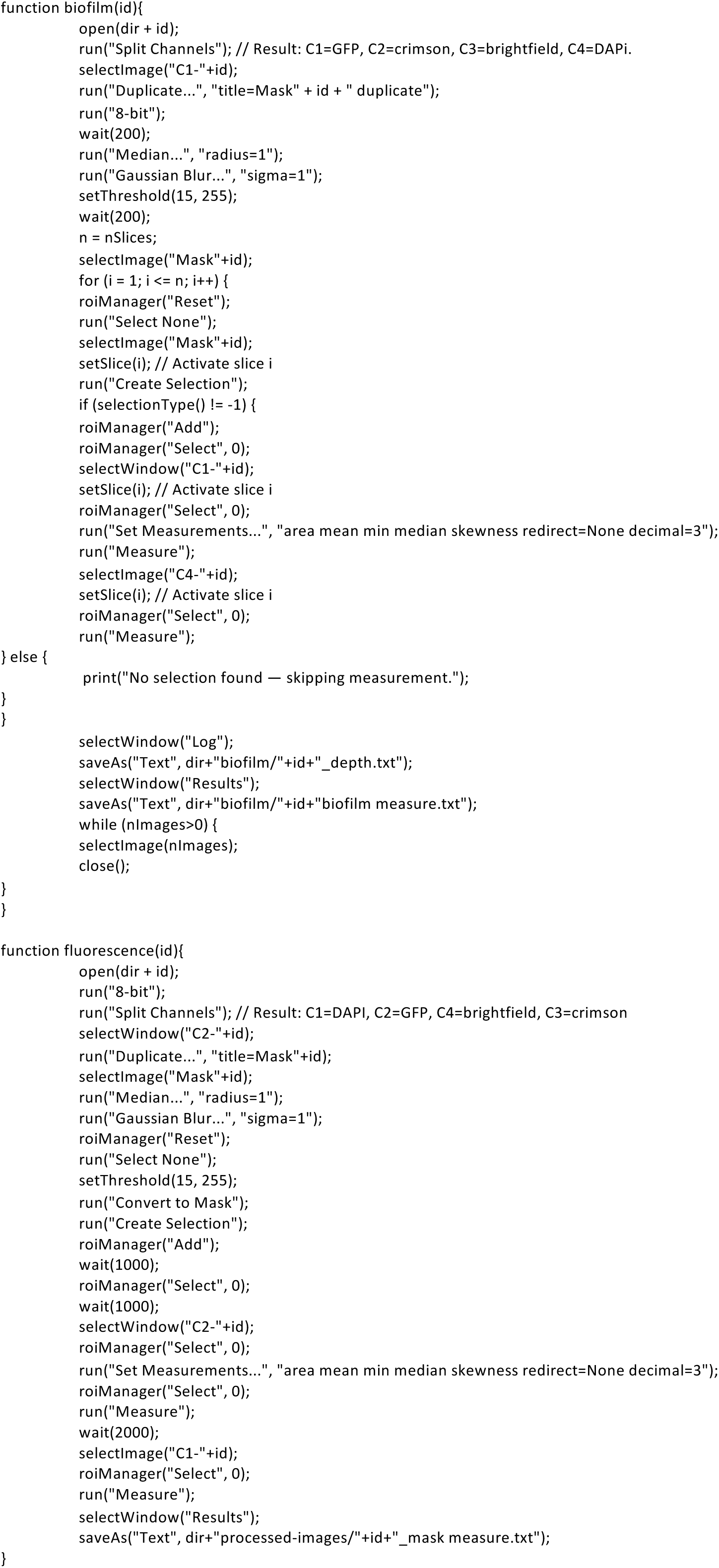

